# The alternative reality of plant mitochondrial DNA: one ring does not rule them all

**DOI:** 10.1101/564278

**Authors:** Alexander Kozik, Beth A. Rowan, Dean Lavelle, Lidija Berke, M. Eric Schranz, Richard W. Michelmore, Alan C. Christensen

## Abstract

Plant mitochondrial genomes are usually assembled and displayed as circular maps based on the widely-held view across the broad community of life scientists that circular genome-sized molecules are the primary form of plant mitochondrial DNA, despite the understanding by plant mitochondrial researchers that this is an inaccurate and outdated concept. Many plant mitochondrial genomes have one or more pairs of large repeats that can act as sites for inter- or intramolecular recombination, leading to multiple alternative genomic arrangements (isoforms). Most mitochondrial genomes have been assembled using methods that were unable to capture the complete spectrum of isoforms within a species, leading to an incomplete inference of their structure and recombinational activity. To document and investigate underlying reasons for structural diversity in plant mitochondrial DNA, we used long-read (PacBio) and short-read (Illumina) sequencing data to assemble and compare mitochondrial genomes of domesticated (*Lactuca sativa*) and wild (*L. saligna* and *L. serriola*) lettuce species. This allowed us to characterize a comprehensive, complex set of isoforms within each species and compare genome structures between species. Physical analysis of *L. sativa* mtDNA molecules by fluorescence microscopy revealed a variety of linear, branched linear, and circular structures. The mitochondrial genomes for *L. sativa* and *L. serriola* were identical in sequence and arrangement and differed substantially from *L. saligna,* indicating that the mitochondrial genome structure did not change during domestication. From the isoforms evident in our data, we inferred that recombination occurs at repeats of all sizes at variable frequencies. The differences in genome structure between *L. saligna* and the two other *Lactuca* species can be largely explained by rare recombination events that rearranged the structure. Our data demonstrate that representations of plant mitochondrial DNA as simple, genome-sized circular molecules are not accurate descriptions of their true nature and that in reality plant mitochondrial DNA is a complex, dynamic mixture of forms.

**AUTHOR SUMMARY:** Plant mitochondrial genomes are commonly depicted in research articles and textbooks as circular molecules that are the size of the genome. Although research on mitochondrial DNA (mtDNA) over the past few decades has revealed that genome-sized circles are exceedingly rare and that alternative forms of mtDNA are more common, many biologists still perceive circular maps as representing one or more physical chromosomes. This misconception can lead to biases in how mitochondrial genomes are assembled and misinterpretation of their evolutionary relationships, synteny, and histories. In this study, we present an assembly methodology that uses short- and long-read sequencing data to determine the mitochondrial genome structures of three lettuce species. We show that these mitochondrial genomes are fluid and dynamic, with multiple sequence arrangements of the genome coexisting within individuals of the same species. Differences in sequence arrangements between species can be explained by rare recombination events. Inspection of physical molecules of mtDNA reveals primarily non-circular forms. We demonstrate that plant mitochondrial genomes are a complex mixture of physical forms and sequence arrangements. Our data suggest that plant mitochondrial genomes should be presented as multiple sequence units showing their variable and dynamic connections, rather than as circles.

## INTRODUCTION

Unlike the relatively simple mitochondrial genomes of animals, the genomes of non-parasitic flowering plant mitochondria are large and complex (1–8). They exhibit extensive variation in size (191 kb – 11,319 kb), sequence arrangement, and repeat content, yet coding sequences are highly conserved (typically 24 core genes with 17 variable genes) (9–13). Mitochondria in plants not only have important roles in respiration, metabolism, and programmed cell death (similar to animal mitochondria), but also in conferring male sterility (14–17). Their evolution has been the subject of numerous studies (3,9,10,18–25).

Because they can often be assembled and mapped as circles, there is a commonly-held misconception that plant mitochondrial genomes exist *in vivo* as circular molecules (the master circle model (26–28)). This was the consensus view until a lack of strong evidence for genome-sized circular molecules in plants and accumulating evidence for non-circular forms caused a shift in the understanding among domain experts and a transition to a more accurate understanding of plant mtDNA as primarily a collection of dynamic non-circular forms (5,6,29–33). These forms can change during development and in response to stress, such as prolonged exposure to low temperatures (34). However, most biologists outside of the specialized community of plant mitochondrial researchers still hold the outdated “master circle” view. This is perhaps because many contemporary publications of mitochondrial genomes persist in presenting a “master circle,” often without mention of any other forms (S1 Table) and this is what is currently presented in most biology textbooks. Additionally, the replication and recombination mechanisms of plant mitochondrial genomes are still not fully understood, nor are the adaptive reasons for the striking differences from animal mitochondrial genomes. Accurate characterizations of plant mitochondrial genome structures are required for understanding their functions, replication, inheritance, and their peculiar evolutionary trajectories.

DNA sequencing and comparisons between sister taxa have revealed that plant mitochondrial synonymous mutation rates are substantially lower than in animal mitochondria, or the plant nucleus (35–38). In contrast, once entire genomes were sequenced, it became clear that while there was conservation of gene content, there was very little conservation of gene order even between close relatives (39, 40), likely due to frequent DNA repair that occurs via homologous recombination and nonhomologous end-joining (22,25,41). Most plant mitochondrial genomes include a substantial fraction of poorly conserved DNA of unknown function and some have dramatically increased their genome sizes to millions of nucleotides, while still encoding only dozens of genes (11,42,43). In contrast to their representation as master circles, mtDNA exhibits complex and dynamic structures, including linear and branched molecules (which could be intermediates in replication or recombination) and these may represent multiple isoforms of the genome (44).

Plant mitochondrial genomes generally have a small number of non-tandem direct or inverted repeat sequences of several kb in length. These may recombine frequently and symmetrically, isomerizing the genome (45, 46). Some genomes assemble into more than one independent molecule (47), although these may also be the consequences of assembly methods and parameters used during the assembly process (11, 48). In many cases, particularly those sequences assembled solely from short-read sequencing, recombination and isomerization have simply been assumed or ignored. There are also dispersed repeats of up to a few hundred base pairs that recombine at relatively low frequencies in wild type plants but often recombine more frequently (and asymmetrically) in DNA maintenance and repair mutants (49–53). Such repeats are not always annotated, but their important contribution to the rearrangement and evolution of mitochondrial genomes is starting to emerge (22).

We employed new technologies to address the challenge of assembling genomes with branches and rearrangements by determining the complete mitochondrial genome sequences of three closely related species in the genus *Lactuca*. In terms of gene content, overall size, and repeat content, these genomes are typical of many flowering plant mitochondrial genomes (24, 25). We combined long-read and high coverage, short-read data to determine the sequences, junctions, rearrangements, and stoichiometry with great precision to produce the first high-quality mitochondrial assemblies with detailed information about structures and isoforms for species in the Compositae. This allowed us to evaluate the repeat structure and frequent isomerization by recombination at the large repeats. We propose a model for rare recombination events that rearranged the mitochondrial genome during the divergence of two *Lactuca* lineages. The physical structure of the genomes visualized by fluorescence microscopy showed that the mtDNA of *L. sativa* exists primarily in branched, linear forms, and subgenome-sized circles. Our data allowed us to document the diversity of isoforms in great detail, clarify misconceptions about mtDNA structures, explore the best methods for assembly of these dynamic, complex genomes, and examine the evolution of mitochondrial genomes in both a wild and a domesticated descendent from a common *Lactuca* ancestor.

## RESULTS

### Overview of assembly of the mitochondrial genomes of three lettuce species

We sequenced and assembled the mitochondrial genomes of three *Lactuca* species: *L. sativa*, *L. serriola*, and *L. saligna.* Because plant mitochondria are known to have dynamic and rearranging genome structures, we recognized the possibility of assembling large contigs with multiple connections to other contigs. Therefore, our assembly approach did not make assumptions about linearity or circularity. We only relied on the sequence reads to identify the contiguous segments and how they are connected. After producing and polishing contigs to form the primary structural units of the mitochondrial genome, we used junction information to join primary structural units together to form secondary building blocks. Finally, we used the stoichiometry of these blocks to find repeated elements that recombine to generate different isoforms of the genome (S1 Fig).

For *L. sativa* and *L. saligna*, we verified our high quality assemblies using a multistep approach involving processing and analysis of data from several sources (Table 1, S1 Fig). For *L. serriola,* we compared Illumina paired-end and mate-pair reads to the *L. sativa* assembly to determine its mitochondrial genome structure. Because the mitochondrial genome of *L. serriola* was essentially identical to that of *L. sativa*, we focused the majority of our subsequent analyses on comparing *L. saligna* and *L. sativa*.

**Table 1:** Types of read data used for mitochondrial genome assembly and analysis

|  | Illumina paired-end | Illumina mate-pair | PacBio | Hi-C |
| --- | --- | --- | --- | --- |
| *Lactuca sativa* | ✓ | ✓ | ✓ | ✓ |
| *Lactuca serriola* | ✓ | ✓ |  |  |
| *Lactuca saligna* | ✓ |  | ✓ |  |

### Initial assembly of PacBio reads for *L. sativa* and *L. saligna*

For *L. sativa* and *L. saligna*, we used the CLC Genomics Workbench with Genome Finishing Module to assemble each mitochondrial genome from PacBio mitochondrial reads (selected from whole genome read data). After polishing with Illumina paired-end reads, this process resulted in 11 contigs for *L. sativa* and 10 for *L. saligna* (Fig 1A). Distinct contigs are designated by letters from K to Z and will be referenced in the text accordingly (see contig naming convention in Materials and Methods). The total assembly length of the polished non-redundant contigs (primary structural units; see below) was 314,646 bp for *L. sativa* and 323,254 bp for *L. saligna* (Table 2). Eight contigs (K, L, M, T, R, P, Q, and Z) were very similar between the two species (ranging from 99.8% to nearly 100% similarity at the nucleotide level), although there were minor variations around the contig termini. In *L. saligna*, the contig designated UV was a single contig that contained sequences found in U and W of *L. sativa*. We therefore split this contig into two parts (U and V) to simplify and clarify the visualization and interpretation of contig relationships. Upon refining contig sequences we defined them as the primary structural units of the mitochondrial genome and determined their junctions, relative orientations, and copy number. Mitochondrial genome coordinates of primary structural units on representative GenBank files are shown in S2 Table.

**Fig 1.**
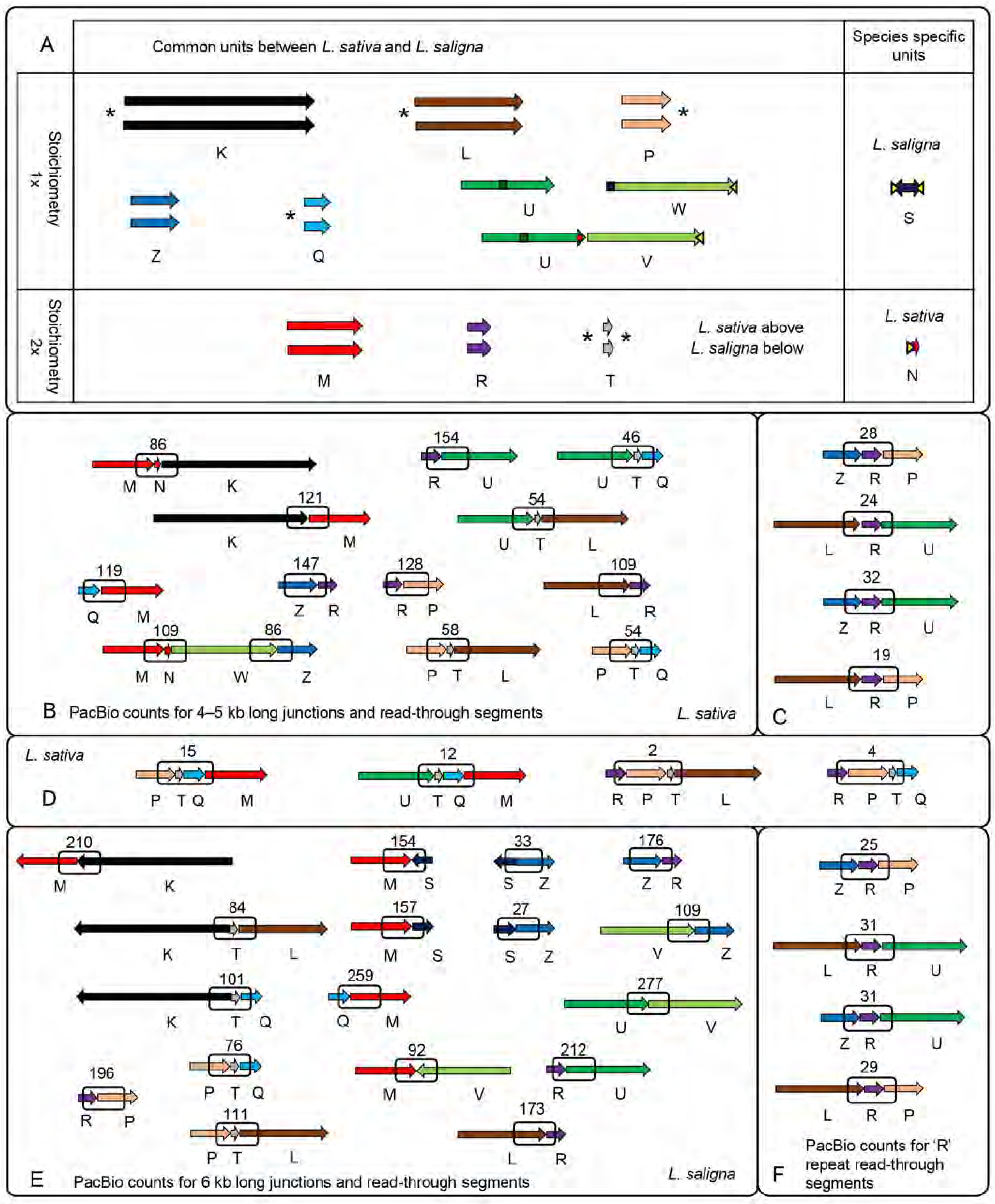
Primary structural units of the Lactuca mitochondrial genome and secondary building blocks. A) Comparison of primary structural units between the *L. sativa* and *L. saligna* mitochondrial genomes. Polished contigs generated from CLC assembly form the primary structural units of the mitochondrial genome and are shown in pairs with *L. sativa* on top and L. saligna on the bottom. The stoichiometry (1x or 2x) and whether the units were common or unique between *L. sativa* and *L. saligna* are indicated. Different termini of a pair of contigs are labeled with an asterisk. The differences between termini can be up to several hundred nucleotides. A dark green bar indicates a large insertion from the chloroplast genome. The contig from *L. saligna*, UV, was a single contig that correspond to major parts of contigs U and W of *L. sativa* and was conceptually split into two segments (color coding) to simplify and clarify the visualization and interpretation of contig relationships. A segment of the S inverted repeat sequence is duplicated in both species on the termini of units W and V, and in unit N of *L. sativa* (yellow arrows). B-F) Secondary building blocks of mitochondrial genomes with a defined sequential order of primary units determined by reverse read mapping (see Materials and Methods) for *L. sativa* (B-D) and *L. saligna* (E,F). Junctions within boxes were quantified using PacBio reads and the counts are displayed as numbers above the boxes.

**Table 2:** Lengths of primary structural units for **L. sativa** and **L. saligna**

| *L. sativa* |  | *L. saligna* |  |
| --- | --- | --- | --- |
| Unit name | Length (bp) | Unit name | Length (bp) |
| K | 78,272 | K | 79,109 |
| L | 44,054 | L | 43,583 |
| W | 53,983 | UV | 90,023 |
| U | 38,613 |  |  |
| M | 30,580 | M | 30,534 |
| P | 20,468 | P | 20,420 |
| Z | 19,295 | Z | 19,278 |
| Q | 11,283 | Q | 11,083 |
| R | 10,430 | R | 10,433 |
|  |  | S | 14,743 |
| N | 4,116 |  |  |
| T | 3,552 | T | 4,048 |
| **TOTAL** | **314,646** | **TOTAL** | **323,254** |

### Classification of repeated sequences

Because segments of the genome that are several kb in length are commonly present at two locations in many mitochondrial genomes (7,29,38–41), we expected that any contigs that represent a segment of the genome present in more than one location would be detected as variation in the copy number of that contig. Indeed, our coverage analysis (S2 Fig) revealed that three contigs (M, R, and T) had twice the coverage relative to the others; thus, we designated these as large repeats. Repeat pairs of M, R, and T would be expected to recombine continuously and serve essentially as hinges between different isoforms of the genome. We also found intermediate-length (< 1 kb) repeats within contigs (S2 Table and GenBank files). For example, a 576 bp repeat that we termed X-01b (yellow triangles on Fig 1) was found in contig N in **L. sativa** and in W/V in both species. In **L. saligna**, this repeat was also present in an inverted orientation at the termini of contig S.

### Identification and characterization of primary structural units and secondary building blocks

To infer the structural topology of the mitochondrial genomes of both species, we took advantage of the long PacBio reads to identify the junctions between contigs. We first polished the contigs by correcting minor discrepancies at their termini and precisely defined their boundaries at junction points. After this refinement, we considered the polished contigs as “primary structural units.” Using the set of maximally-informative reads that we selected during the assembly process (see Materials and Methods), we employed a “reverse read mapping” approach for determining which primary units were joined to one another. In this approach, the primary structural units were fragmented into 2 kb sliding windows with a 1 kb step size and mapped against the maximally-informative read set for each species (see Materials and Methods). This enabled the identification of secondary building blocks composed of several primary structural units in a specific order (Fig 1 B-F). In many instances, the PacBio reads were long enough to span junctions on both sides of the T and R repeat units (but not the much longer M repeat unit). These basic building blocks revealed the existence of several different arrangements of the primary structural units. For example, the 3’ end of unit R was joined to unit U in one building block and unit P in another in both species. Ultra-long reads that spanned junctions between four units (Fig 1D) confirmed these alternative arrangements for **L. sativa**. After re-assembling the junctions for greater accuracy (see Methods), we mapped 2 kb segments spanning the junctions to PacBio reads to quantify how often they occurred (S3 Fig).

### Identification of major isoforms

Given the set of all detected secondary building blocks and their stoichiometry, we were able to order the primary structural units into sequences that were the total length of the mitochondrial genome. In both species, there were several different arrangements (isoforms) of primary structural units that were well supported by the data. The two major mitochondrial isoforms for **L. sativa** can be represented as in Fig 2A. The difference between the two major isoforms (α and β) of **L. sativa** can be described as resulting from an exchange of two primary structural units, P and U, between long repeats R and T. These major isoforms have roughly equivalent stoichiometry. With two copies of each of the three large repeats included, these isoforms represent genome lengths of 363,324 bp for **L. sativa** and 368,269 bp for **L. saligna**.

**Fig 2.**
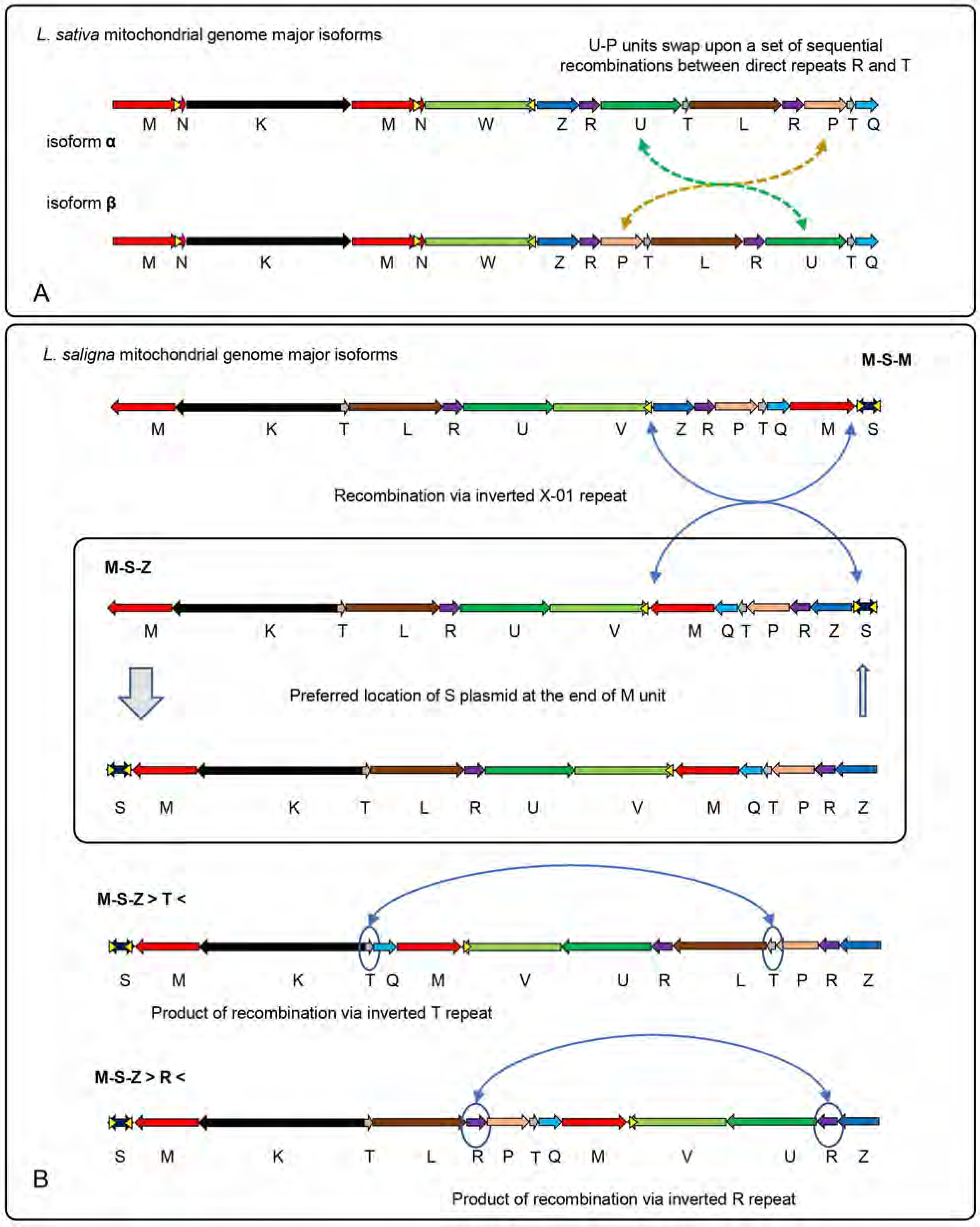
Isoforms of *Lactuca* mitochondrial genomes. Putative major isoforms of *L. sativa* (A) and *L. saligna* (B) were derived from analysis of secondary building blocks (Fig 1 B-F) and the primary structural unit stoichiometry (Fig 1A and S2 Fig). For *L. sativa*, the dashed green arrow in A shows rearrangement of unit U and the dashed yellow arrow shows rearrangement of unit P between the two major isoforms. For *L. saligna*, putative recombination events that result in a transition from one isoform to another are indicated by solid blue arrows. Isoforms are labeled by the configuration of the units M, S, and Z, with > and < symbols designating where recombination has taken place. Yellow triangles indicate repeat X-01. Displayed isoforms are simplified models that fit the sequential order of primary structural units and stoichiometry data and make no assumptions about the underlying form of mtDNA molecules.

The assembly of **L. saligna** was more complex. We identified an interesting 12 kb primary unit in **L. saligna**. This sequence, unit S, includes a Type B2 superfamily DNA polymerase, a T3/T7 phage type RNA polymerase, and a 1,218 bp inverted repeat at each end, resembling linear plasmids described in other plants (54). Some plant mitochondria include integrated fragments of these plasmids or autonomous plasmid molecules (55–60); phylogenetic analysis suggests occasional horizontal transfer of these as well (61). Unit S appears to be an integrated copy of a molecule of this type in 1. **L. saligna**. A portion of the sequence in unit S is present in **L. sativa**, but it lacks one inverted repeat and the DNA and RNA polymerase genes, and is presumably degraded and nonfunctional. Other species that carry such plasmids include relatives of *Lactuca*, *Daucus carota* (59) and *Diplostephium hartwegii* (62). We found no evidence for free linear plasmids in either *Lactuca* species. Unit S is equally likely in either orientation in *L. saligna*, indicating that the inverted repeats at the end recombine with each other at a high frequency.

Fig 2B displays a comprehensive set of **L. saligna** mitochondrial isoforms representing all of the detected secondary building blocks (Fig 1E,F). The presence of the integrated complete S unit in **L. saligna** prevented us from unambiguously constructing a simple and stoichiometrically symmetrical genome model as in **L. sativa**. Primary unit T, which is a repeat, was present in four arrangements: P-T-Q, P-T-L, K-T-L, and K-T-Q. Likewise, the repeat unit R was found in four distinct blocks: Z-R-P, Z-R-U, L-R-U, and L-R-P. Analysis of ultra-long PacBio reads (Fig 1F) suggested that all configurations with the R and T repeats are equally represented (as was also the case for **L. sativa**). Primary unit S was flanked by unit M on both sides and can potentially form an inverted repeat configuration of ∼32 kb. Recombination via the X-01a repeat, which is part of the inverted S repeat structure, and its counterpart in the V unit can lead to the formation of an M-V junction (Fig 2B). PacBio read-through analysis showed the M-V and V-Z junctions in equilibrium (equally represented). There was a noticeable deficiency of S-Z junctions in comparison with M-S junctions, which may reflect that a linear genome structure with unit S at one terminus was more prevalent. These **L. saligna** isoforms were not necessarily in stoichiometric equilibrium (like the major **L. sativa** isoforms) due to anomalies caused by the integrated linear S plasmid.

### Annotation of the sequences

Because the representation of the annotations on circular maps has led to the misperception of the existence of a specific circular molecule, we present annotations of each primary unit separately. Importantly, we do not suggest that these maps represent specific linear chromosomes either. Combining the junction data and the physical data (below) suggests a fluid and dynamic genome with multiple isoforms and topologies that are larger than the primary units. The annotation maps shown in Fig 3 are an effective way to provide a static figure representing the dynamic reality of mitochondrial genomes. Arbitrarily choosing to present one possible isoform among many (which unfortunately is currently required for GenBank submission) leads to an overly simplistic picture of the genome.

**Fig 3.**
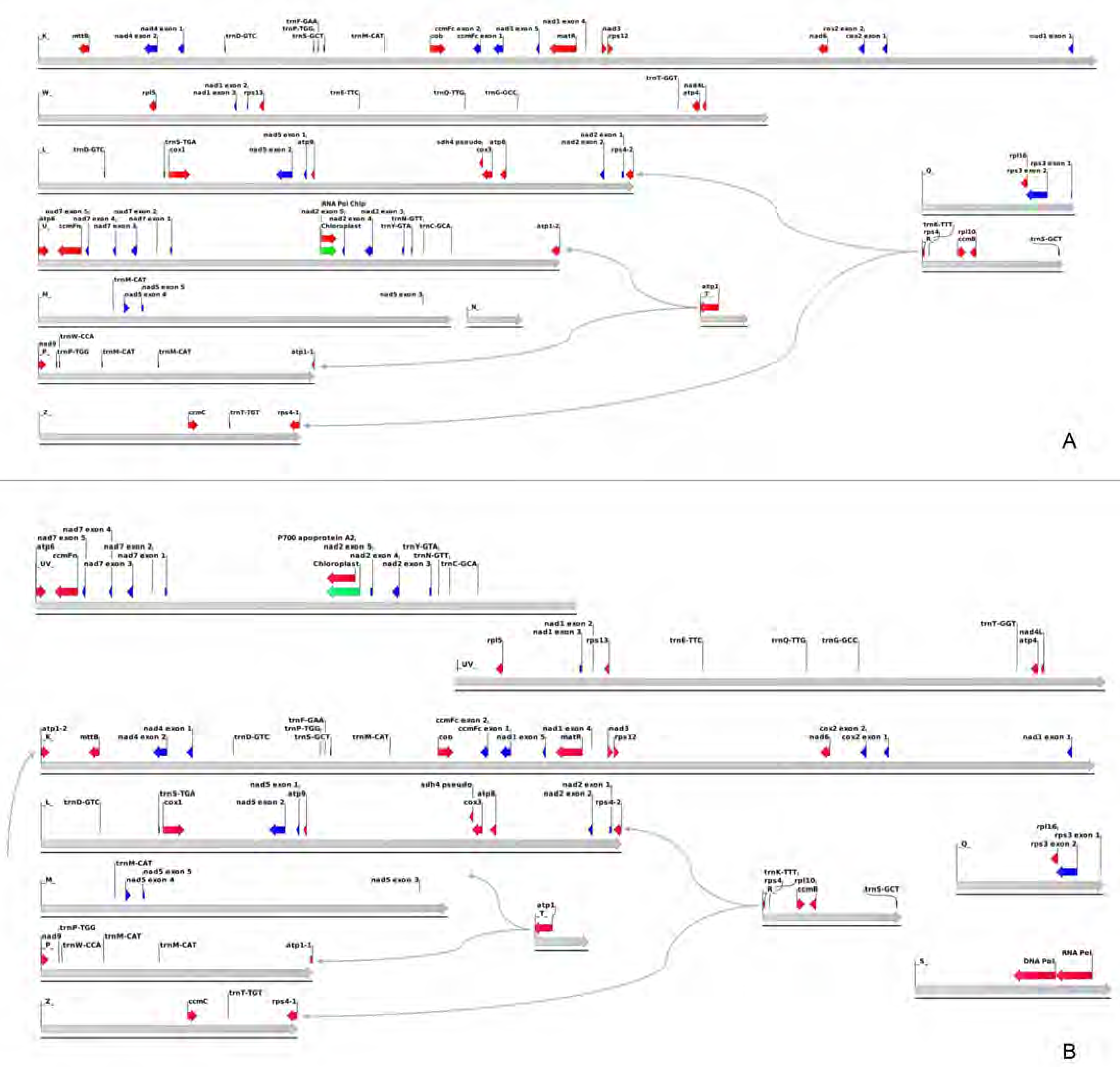
Lactuca mitochondrial genome annotations. The annotations for genes and other sequence features for *L. sativa* and *L. saligna* are displayed along the primary structural units (see Fig 1A) for each genome, which are indicated by thick gray arrows. Intronless genes are indicated by red arrows, exons of spliced genes are indicated by blue arrows, and plastid insertions are indicated by green arrows. Genes that span junctions between primary units are indicated by a jagged line at the division point. Thin gray arrows show alternative junctions between primary structural units that result in different models for the genes that are split over a junction.

Annotation of the genomes identified a set of rRNA, tRNA, and protein-coding genes that is typical of angiosperms. Both **L. sativa** and **L. saligna** have all of the core genes (as defined in reference (12): *atp1*, *atp4*, *atp6*, *atp9*, *ccmB*, *ccmC*, *ccmFc*, *ccmFn*, *cob*, *cox1*, *cox2*, *cox3*, *matR*, *mttB*, *nad1*, *nad2*, *nad3*, *nad4*, *nad4L*, *nad5*, *nad6*, *nad7*, and *nad9*. Of the variable genes defined previously (12), both *Lactuca* genomes have the same gene content as *Helianthus annuus*, namely, *rpl5*, *rpl10*, *rpl16*, *rps1*, *rps3*, *rps4*, *rps12*, and *rps13*. **L. sativa**, **L. saligna**, and *H. annuus* also have a pseudogene of *sdh4*, identified by a stop codon in the middle. This gene content is unremarkable for a member of the Compositae, except for four duplicated genes described below.

The repeat units in *Lactuca* mitochondrial genomes contain protein-coding genes, as is often the case in plant mitochondrial genomes, although the particular genes duplicated varies. The *ccmB* and *rpl10* genes are both located in repeat unit R, resulting in two identical copies in the full-length genomes of both species. Another gene present in two locations is *atp1*, encoding the alpha subunit of the F_o_F_1_ ATP synthase (atpα). Nearly the entire gene is in repeat unit T, including the 5’ flanking region, the start of translation, and 1,319 nucleotides of coding sequence. In flowering plants, the *atp1* gene encodes a highly conserved protein of between 505 and 512 amino acids in length. One copy in both *Lactuca* species, annotated as *atp1-1*, extends from repeat unit T into unit P and encodes a 511 amino acid protein that is 99.8% identical to the atpα proteins from *H. annuus* and *D. hartwegii* (62, 63) and is 97.3% identical to the atpα protein of the distantly related dicot *Vitis vinifera* (64). The second copy, *atp1-2*, extends from repeat unit T into unit U in **L. sativa** and into unit K in **L. saligna**. These copies are identical to *atp1-1* through 444 amino acids of coding sequence and then diverge: *atp1-2* of **L. sativa** adds an additional 205 amino acids and *atp1-2* of **L. saligna** adds 216. A similar situation exists in *D. carota* (65) in which the first 1,452 nucleotides of the gene are present in a repeat. One copy (annotated in (65) as a pseudogene) is a 514 amino acid protein that resembles other plant atpα proteins, and the other, annotated as *atp1*, includes the first 485 amino acids of atpα fused to an additional 271 amino acids, suggesting that the *D. carota* annotations are reversed. Similarly, the 5’ end of *rps4* is in repeat unit R, joined to units Z and L (in both *Lactuca* species). The gene that extends from R into Z is annotated as *rps4-1* and resembles other plant mitochondrial *rps4* genes. The other copy, *rps4-2*, consists of 189 nucleotides encoding the first 63 amino acids of the rps4 protein, fused to a 268 amino acid open reading frame in unit L, which does not resemble other plant rps4 proteins. Further work is needed to understand if these novel chimeras are expressed, whether they are selectively neutral or have a novel function, and how assembly of multi-subunit protein complexes is regulated.

### Long distance sequence technologies support mitochondrial genome isoform models

Illumina mate-pair and Hi-C libraries validated the models of the mitochondrial genome generated using PacBio reads. Reads from two different insert size libraries (2.5 and 10 kb) for **L. sativa** and *L. serriola* mapped to the mitochondrial genome isoform alpha confirmed the sequential order of the primary structural units. The distribution of long repeats (R, T, and MN) on two-dimensional distance plots were consistent with the structure of isoforms inferred from the reverse mapping approach using PacBio reads (Fig 4A,B). The distribution of long-distance interactions revealed by mapping reads from the Hi-C genomic library of **L. sativa** to the mitochondrial reference assembly was congruent with mate-pair analysis (Fig 4C). Mitochondrial DNA is not naked *in vivo*, but is complexed with proteins into a higher order structure known as the nucleoid (66–68). These structures are not well-characterized but are very likely to be important for mitochondrial genome replication, inheritance, and transcription (68–71). Hi-C has the potential to detect contacts between regions of the genome that are brought into proximity *in vivo* by proteins or other components of the mitochondrial nucleoid, but we did not find strong evidence for long-distance interactions arising from the organization of mtDNA within the nucleoid (see also S4 Fig).

**Fig 4.**
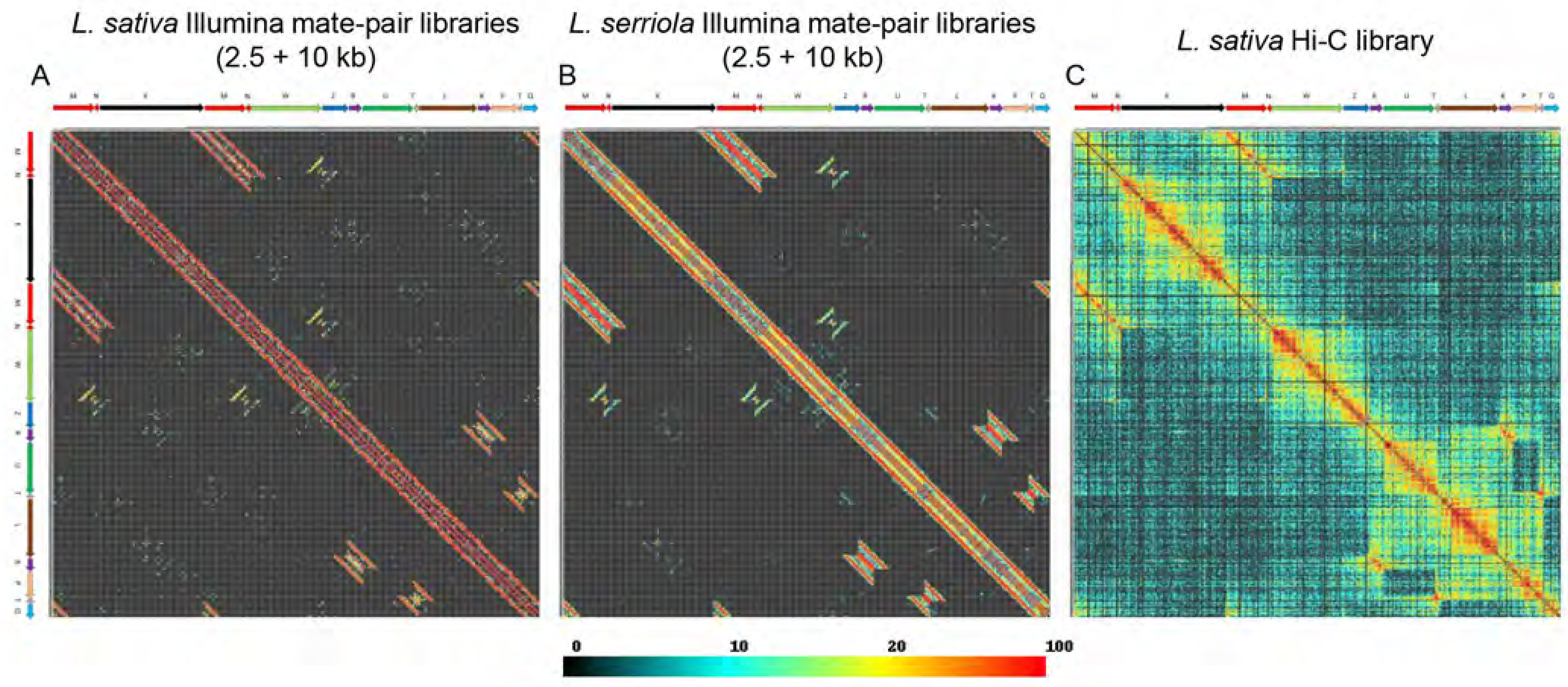
Long distance analysis of mitochondrial genomes using Illumina mate pair and Hi-C libraries. Plots of distances between read-pairs Illumina mate pair libraries with 2.5 and 10 kb insert sizes in 1 kb bins for *L. sativa* (A) and L. serriola (B) exhibit essentially identical long-distance sequence connections. C) Plot of Hi-C contact frequencies for *L. sativa*. Major isoform α was arbitrarily chosen as a reference for visualization of mate-pair distances and Hi-C contact frequency. The color gradient displayed below panels reflects the number of read-pairs (out of a total of 320,000) in each 2-dimensional-1-kb bin and applies to all panels.

Detailed inspection of distance plots from Illumina mate-pair reads for junction regions revealed the existence of minor isoforms that resulted from recombination between short repeats of less than 600 nt (Fig 5). Presence of these minor isoforms did not affect the equilibrium between the major isoforms of **L. sativa** (Fig 2A). Plots shown in Figs 4 and 5 were generated by mapping 320,000 mitochondrial reads each for 2.5 and 10 kb libraries. To detect recombination between short repeats of less than 300 nt, all available (∼1 million) reads from the 5 and 10 kb mate-pair libraries for **L. sativa** and the 10 kb library for *L. serriola* were used. To simplify the analysis of recombination at short repeats, which can be ambiguous and complicated due to reads aligning at multiple locations, only unique mappings were selected (S5A-C Fig). All mappings of reads from **L. sativa** and *L. serriola* generated similar patterns with detectable signals of recombination events between short repeats. The fraction of recombinants detected between short repeats was very low and estimated to be within (1% – 10% for repeats 100 – 500 bp and less than 1% for repeats shorter than 100 bp) of the values on the main diagonal (see example data in S5D Fig). This rarity explained why only a few corresponding recombinants were detected within the PacBio reads.

**Fig 5.**
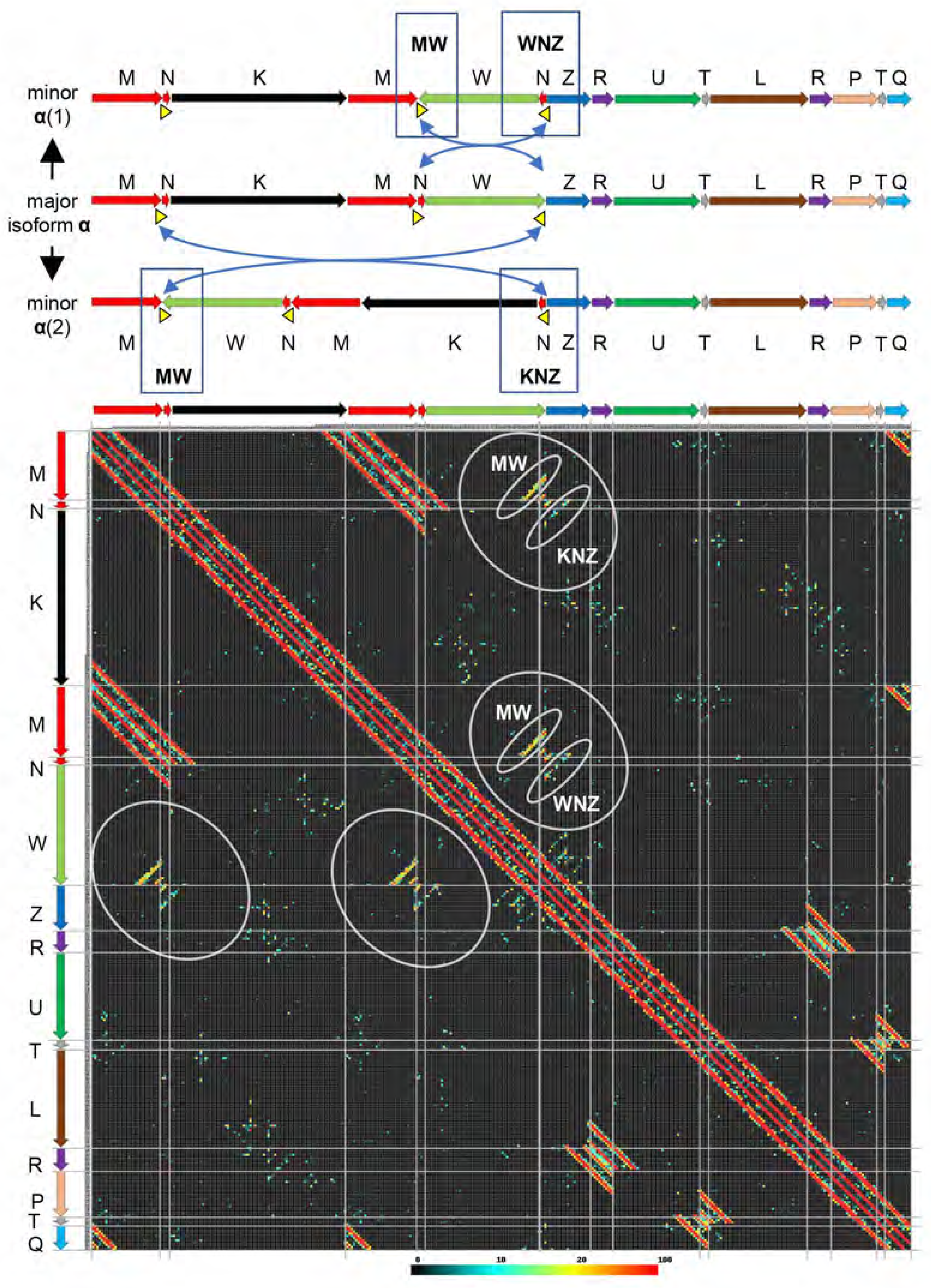
Detection of minor isoforms using Illumina mate pair data. Above, schematic illustrations of the transition between major isoform α and two minor isoforms through recombination at repeat X-01b (yellow triangles and blue arrows). Below, high-resolution-Illumina-mate-pair plot showing sequence connections (indicated by gray ovals) that are expected for the minor isoforms shown above. Mate-pair library sizes were 2.5 and 10 kb. The major mitochondrial isoform of *L. sativa* includes the following order of basic units: M-N-K-M-N-W-Z. There is a medium size repeat X-01b of length ∼500 bp (yellow triangle) at the N and W termini that can cause two distinct recombination events as shown on the right side of the figure. Upon X-01b recombination, two new isoforms have three distinct junctions M-W (common for both), W-N-Z, and K-N-Z (specific for each isoform). A long distance plot of mate-pair libraries clearly demonstrated the existence of both isoforms and all three junctions. Ovals highlight the diagonals of long distance interactions that were evidence of the existence of minor isoforms for the *L. sativa* mitochondrial genome. Similar patterns were detected using L. serriola mate pair libraries (Fig 4B and S5C Fig). The color gradient displayed below panels reflects the total number of read-pairs (out of a total of 320,000) in each 2-dimensional-1-kb bin and applies to all panels.

### Evolution of the mitochondrial genomes of **L. sativa** and **L. saligna**

While **L. sativa** and *L. serriola* are identical in their sequence arrangement and repeat content, **L. saligna** has notable differences from **L. sativa**. A major difference in the **L. saligna** lineage is that there has been an inversion around the intermediate-sized repeat X-07. Conceptual reversal of that inversion leads to an intermediate form that can be converted to **L. sativa** by additional rearrangements (Fig 6). Without an outgroup or intermediates in the evolutionary process, it is not currently possible to unambiguously specify the molecular events in each lineage that led to these differences. An additional important difference is the 12-kb sequence unit S integrated in the genome of **L. saligna** and partially deleted in **L. sativa**.

**Fig 6.**
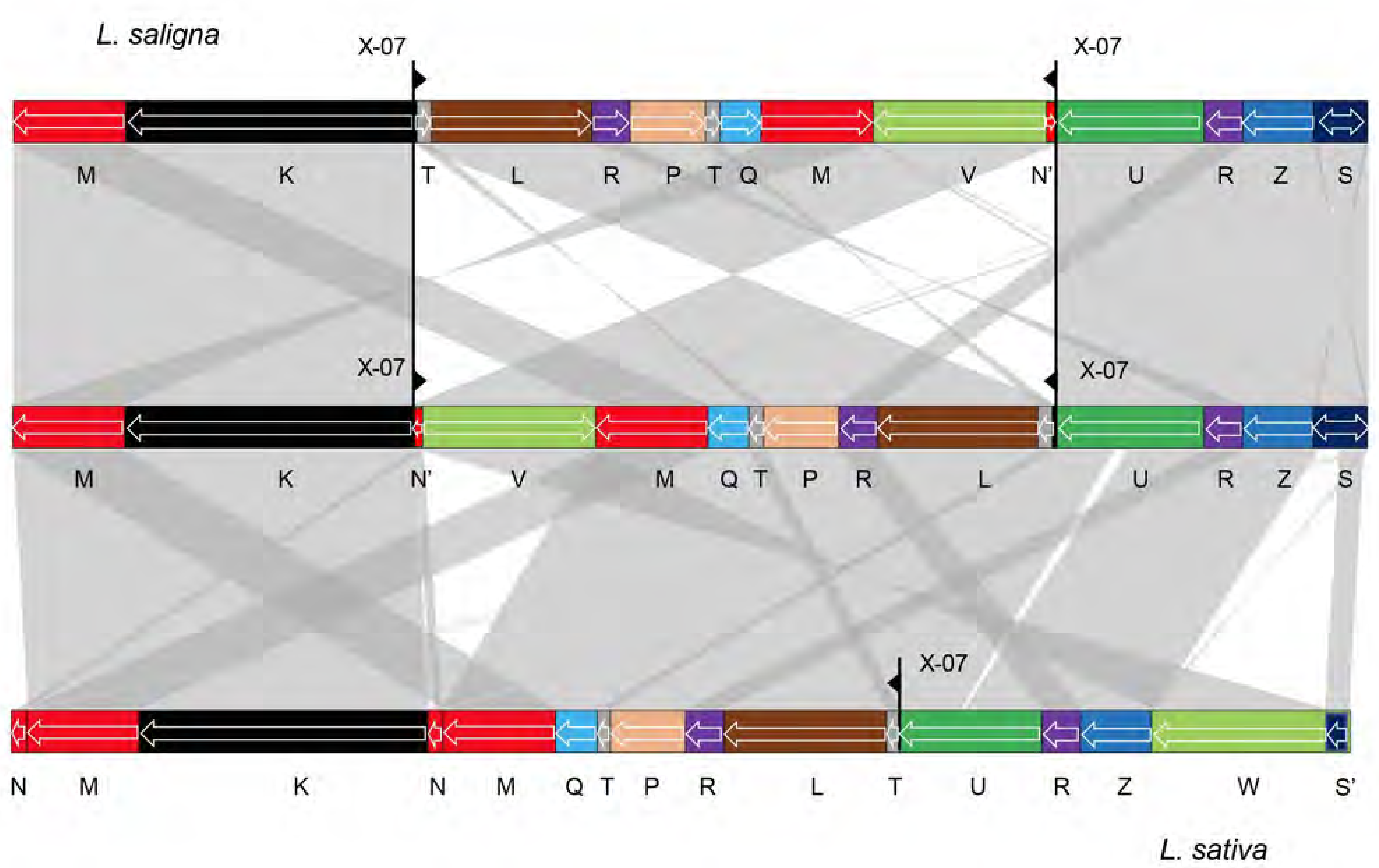
Interconversion between *L. sativa* and *L. saligna* genomes. Linear representation of one of the *L. saligna* major mitochondrial genome isoforms (GenBank accession MK759657) on the top. The middle structure is a putative minor isoform of *L. saligna* derived from recombination at repeat X-07 (black triangles), which leads to a similar arrangement of primary structural units as that of *L. sativa* major isoform α at bottom (GenBank accession MK642355). Collinear segments (longer than 500 bp) are indicated by gray shading.

In addition to the inversion and the integrated linear plasmid, there is also a complex rearrangement and a number of minor differences distinguishing the two *Lactuca* lineages. Some of these differences may be due to events that began with a homologous strand invasion, but included additional breakage events in the unique flanking regions that were subsequently repaired by non-homology based mechanisms. Footprints of such events include a 472 bp sequence at the end of **L. sativa** unit L found at the end of **L. saligna** unit T, and a short segment of ∼200 bp at one end of **L. sativa** unit Q missing in **L. saligna**. In addition, recombination at repeat X-07 likely explains a ∼600 bp segment located at the end of unit K in **L. saligna** that is instead at the end of unit U in **L. sativa**. Also, **L. sativa** has a distinct unit N, but a small portion of it is present in **L. saligna** unit U. Fragments of the chloroplast genome are integrated into the same location in unit U in both species; remarkably, the integrated sequences are not the same. In **L. sativa*,* the integrated sequence is a fragment of a DNA-directed RNA polymerase gene; in **L. saligna**, it is a fragment of a P700 chlorophyll a apoprotein gene (*psaB*). Side-by-side comparison of mitochondrial genome primary structural units for both species is shown on the diagonal plot in S6 Fig.

### Physical structure of mitochondrial DNA molecules in **L. sativa** confirms multiple forms of the genome

We observed a variety of physical structures of mtDNA molecules using in-gel fluorescence microscopy after staining agarose-embedded mtDNA from **L. sativa** with DNA-binding fluorophores (Fig 7). We quantified the structural forms in multiple microscopic fields (Fig 7A-F), according to the primary form present in the field. Branched linear forms were most frequently observed (40 of 98 fields); circular molecules were observed in only 22 fields (Fig 7G). Additional examples of branched forms are shown in S7 Fig. Length measurements of the 23 circular molecules observed in the 22 fields of circular DNA (Fig 7H) revealed no molecules that were the expected length of the ∼363-kb genome (121 μm), and only smaller circles with a mean length of 37.6 μm (approximately 113 kb). Estimating the length in kb from the length in μm assumes that there is no compaction of the DNA. We conclude that the most likely interpretation of our data is that all of these are subgenomic circles, unless the DNA is heavily compacted. Furthermore, no supercoiled circular or genome-sized linear molecules were observed by pulsed-field gel electrophoresis (Fig 7I) in five different mitochondrial samples from **L. sativa**. The majority of DNA in the gel was confined to the well, consistent with the immobilities of branched linear and relaxed circular forms. The remainder ran as a small smear of linear fragments ranging in size from ∼50 to 100 kb. No band representing the linear DNA form of the mitochondrial genome (isoforms α and β in Fig 2A; ∼363 kb) was observed. These results indicate that simple and branched linear forms are the predominant form of mtDNA molecules in **L. sativa**.

**Fig 7.**
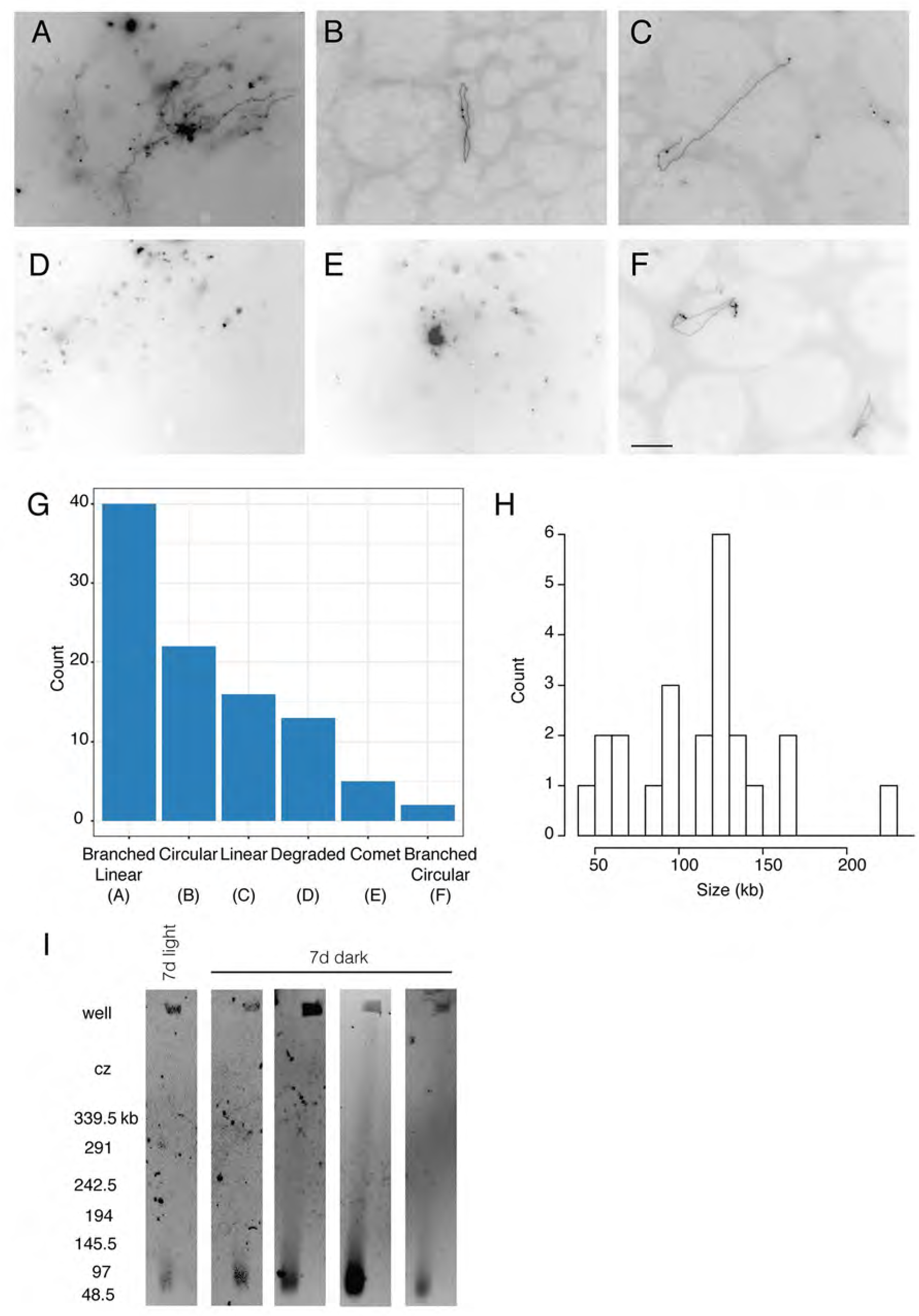
Structural analysis of *L. sativa* mitochondrial DNA by in-gel fluorescence microscopy and pulsed-field gel electrophoresis. (A-F) DNA obtained from mitochondria isolated from seven-day-old dark-grown seedlings (roots and shoots) was stained with either ethidium bromide or QuantiFluor® dye. Images are representative of branched linear (A), circular (B), linear (C), degraded (D), comet (E), and branched circular (F) structures. The scale bar in panel F applies to panels A-F and is 10 μm, corresponding to the length of approximately 30 kb of DNA. Branched linear: interconnected linear forms with or without a densely staining central core. Circular: closed loop without any additional branches. Linear: linear fiber with no branches. Degraded: many small molecules of undetermined structure. Comet: bright core with short connected fibers and no other visible branch points. Branched Circular: closed loop structure with linear branches. G) Quantification of the primary structures observed among 98 total microscopic fields. H) Size distribution of the circular molecules based on length measurements of their circumferences. I) Pulsed-field gel electrophoresis. Each lane represents an independent gel run. Sizes (in kb) are determined from the migration of lambda concatemers run in each gel experiment. In some cases, the DNA migrated at a slight angle to the left.

## DISCUSSION

The complexity of plant mitochondrial genomes has confounded efforts to characterize their sequence, structure, and dynamic evolution. Despite the general acceptance among mitochondrial biologists that plant mitochondrial genomes have a variety of configurations (5,6,29,30,32–34,72), over half of the publications on plant mitochondrial genomes since 2016 have presented circles as if they are the only form or the primary form of the genome, without any indication that the concept of a “master circle” is only a model (see S1 Table). Assumptions have to be made in any genome assembly process about the type of molecule being assembled and published sequences reflect choices made during the assembly process. Because plant mitochondrial genomes rearrange so frequently and are present in so many complex and non-circular states, assumptions as to the circularity of the genome before assembly have inevitably led to arbitrarily circularized structures (73, 74) and incomplete and/or incorrect structures. In addition, GenBank restricts sequences to being designated as either circular or linear, with no provision in the annotations for dynamic structural forms. When plant mitochondrial genomes are assembled with the goal of producing a “master circle,” and their sequences are archived as such, it perpetuates the outdated notion of the existence of master circle molecules, especially for scientists who are outside of the specialized community of plant mitochondrial researchers.

In addition to assuming a master circle, sometimes the goal has been to assemble the sequence into unique circular molecules of minimum size, in spite of the existence of very large repeats in different molecules that would allow for recombination to combine smaller molecules into larger ones (11, 48). Assemblies reported as multi-chromosomal may not exist as distinct molecules. In *Silene conica* (11), for example, the assembled constructs referred to as chromosomes 9, 11, 12, 14, and 23 total 600 bp in length; however, 389 kb of that length consists of repeats between 1 and 75 kb, matching sequences in 13 other assembled circles. These repeats can act as substrates for recombination and combine multiple chromosomes into a single sequence. We suggest that a more comprehensive analysis of structure could inform assembly choices in order to produce a more accurate understanding of plant mitochondrial genomes. Incomplete assemblies may also be missing such repeats that do exist *in vivo* and lead to the incorrect interpretation of multiple molecules. Genome rearrangements can occur both within species or individuals in real time and between species during their evolution and divergence (22); therefore, accurate assembly and identification of multiple isoforms is critical to understanding the evolution of mitochondrial genomes.

In this study, we aimed to produce accurate assemblies and characterize isoforms of the mitochondrial genomes of three *Lactuca* species. Our success was dependent on the exploitation of multiple long-read technologies and atypical assembly approaches. Our use of Illumina mate-pair, Hi-C, and long PacBio reads enabled the high resolution analysis of the mitochondrial genomes of three species of *Lactuca*. Long reads at high coverage depth were essential for accurately determining the *in vivo* sequences and arrangements because short paired-end reads would not allow the identification of recombination events that involve repeats longer than the paired-end library fragments. We assembled these genomes with CLC, reverse read mapping, and long distance analysis with mate-pair reads that made no assumptions as to the component structures and redundancies. Conventional assembly software programs, such as Falcon or Canu (75, 76), assume a single chromosomal sequence and consequently are limited in their ability to identify all isoforms that may exist in plant mitochondrial genomes and resolve complex nested repeats. Assemblers that can split putative chimeras (isoforms) into distinct structural units in the initial steps of assembly have advantages over those that try to assemble all reads into a single linear or circular genome. Bottom-up construction of contigs and alignment to raw PacBio reads allowed a sensitive and accurate assembly of dynamic mitochondrial genomes and identification of multiple isoforms, resulting in precise representation of the subtle complexities of plant mitochondrial genomes and enabling evolutionary studies. The availability of long read sequences for an increasing number of plant species provides the opportunity for the reassessment of mitochondrial structures and diversity across the plant kingdom using our approach. When we applied our workflow to the mitochondrial genome data of *Leucaena trichandra* (74), we were able to generate a cyclic graph for this genome and connect all segments into contiguous isoforms (S8 Fig).

Our structural studies using fluorescence microscopy of mtDNA molecules and Hi-C provided a detailed description of the complexities of plant mitochondrial genome architecture and dynamics that was consistent with our genome assemblies. This provided further evidence for the existence of multiple major and minor isoforms produced by homologous recombination at very large repeats. The prevalence of branched and linear forms of mtDNA likely represents ongoing recombination, perhaps due to recombination-dependent DNA replication (5). The presence of smaller linear and degraded forms might be evidence of post-replication processing. In *Chenopodium album* cell cultures (6) and in later stage mung bean cotyledons (34), the complexity first increases during mtDNA replication and is then processed to simpler forms after replication has been completed. We would expect that the mitochondria isolated from 7-day-old whole lettuce seedlings would be a mix of those in the DNA replication stage and those post-replication and our analysis of the DNA structural forms is consistent with this expectation. **L. sativa** major isoforms had a genome size of ∼363 kb, but no linear band of this size was observed by pulsed-field gel electrophoresis (PFGE). Circles only accounted for a small fraction of the DNA forms, yet the assemblies had a circular topology. This is similar to what has been observed with PFGE of watermelon (77), tobacco (5), soybean (32), and maize (78). Although there was a range of percentages of molecules in each structural category, circles never accounted for the majority of the mtDNA and exhibited a broad distribution. The interpretation that the branched linear structures contain multi-genomic concatemers of the genome is consistent with all of these findings. The Hi-C data also suggest no higher-order structure of mtDNA within the nucleoid.

Repeated sequences in plant mitochondria provide the substrates for contemporaneous and long-term structural variation. Large repeats, usually several kb or more in length, recombine frequently and isomerize the genome continuously. Whether the abundance of isoforms varies among cell types or tissues remains to be investigated; however, the tools are now available for the detailed analysis of specificity. There are also infrequently recombining non-tandem repeats, usually less than several hundred base pairs in length (25). Although it is unclear what controls the interconversion of isoforms and their relative stoichiometries, repeat length may be an important factor. Homology-based ectopic recombination for the intermediate-length repeats has been shown to occur following double-strand breaks (52,79–81), when DNA repair functions are impaired by mutations (49,52,82), and occasionally during the evolution and divergence of species (22). We were able to compare our detailed structures of the different species and infer the most parsimonious path from possible ancestral forms to the genomes of the extant species. Ectopic recombination between dispersed repeats could explain some of the structural changes that have become fixed between the two species. We also identified small linear plasmid-like molecules that have been shown to exist autonomously in plant mitochondria in a variety of species and are sometimes integrated into the mitochondrial genome (reviewed in (83)). Both *Zea mays* and *D. carota* have integrated plasmids in their mitochondrial genomes which prevented the assembly of the genomes into circular maps (65,84,85). One such linear plasmid is integrated into the **L. saligna** genome but is incomplete in **L. sativa**. We found no evidence for its integration at any other location, nor for its existence as an autonomous molecule. Its presence in the **L. saligna** genome and its frequent inversions complicated the analysis of junctions in **L. saligna** and prevented a simple interpretation of the stoichiometry of the junctions.

This is the most detailed description of the sequence, structure, and dynamics of plant mitochondrial genomes to date. Our comprehensive approach for investigating mitochondrial genomes can be applied to existing and future datasets generated for other plant species. This approach should avoid incomplete assemblies and reveal the complexity of multiple isoforms, non-circular molecules within circularly permuted maps, and recombination events that occur within and between species. This will be facilitated by the advent of even longer reads provided by rapidly advancing single molecule (PacBio) and nanopore sequencing (Oxford Nanopore Technologies). Understanding the evolutionary changes that can occur with mitochondrial plant genomes, including integration or loss of linear plasmid-like molecules, provides the foundation for future work on plant evolution and taxonomy, as well as mitochondrial genome structure and function.

## MATERIALS AND METHODS

### Germplasm sources

**L. sativa** cv. Salinas is a designated cultivar and seeds can be obtained from multiple commercial sources. Seeds of the **L. saligna** accession are available from the Centre for Genetic Resources, the Netherlands (https://www.wur.nl/en/Research-Results/Statutory-research-tasks/Centre-for-Genetic-Resources-the-Netherlands-1/Expertise-areas/Plant-Genetic-Resources.htm) under the stock number CGN5271.

We used our UC Davis *L. serriola* stock US96UC23 and will provide seeds upon request.

### PacBio reads

DNA was extracted from seven-day-old dark-grown seedlings of **L. sativa** cv. Salinas grown in sterile conditions at 15°C using a modified CTAB protocol (86) with two chloroform extractions. DNA was removed after EtOH precipitation using a glass hook and then washed two times in 70% EtOH. The DNA was further processed with a high-salt, phenol-chloroform extraction and precipitation and removal of polysaccharides (https://www.pacb.com/wp-content/uploads/2015/09/Experimental-Protocol-Guidelines-for-Using-a-Salt-Chloroform-Wash-to-Clean-Up-gDNA.pdf). Finally, the DNA was precipitated using EtOH and washed two times with 70% EtOH. The DNA was divided into two samples for library preparation. For the first library, the DNA was sheared using a Megaruptor instrument to 20 kb before PacBio library construction, while the second PacBio library was prepared directly from the non-sheared DNA. SMRTbell libraries were made according to manufacturer’s standard protocol (Pacific Biosciences). Libraries were size selected > 20 kb using BluePippin (Sage Science). Sequencing on a Pacific Biosciences (PacBio) RSII Instrument generated 39.6 Gb of 2.7 million single pass reads from 28 SMRT cells. The raw reads were deposited in GenBank under accession numbers SRX3557844–SRX3557871.

For the **L. saligna** germplasm accession CGN5271, DNA was extracted from leaves of young seedlings grown on soil in a greenhouse under long-day (16 h light) conditions with a temperature of 21°C during the day and 19°C at night using a modified version of the protocol described previously (87). PacBio long read data was obtained from 86 SMRT cells. The selected set of mitochondrial reads (1 Gb, 101,753 reads) is available at SRA under accession number SRX5104332.

### Illumina genomic reads

For **L. sativa** cv. Salinas, we used previously published Illumina whole genome libraries (88). Paired-end reads included insert sizes of 175 bp (SRR577192), 475 bp (SRR577183), and 750 bp (SRR577184), and mate pair reads included insert sizes of 2.5 kb (SRR577197), 5 kb (SRR577193), and 10 kb (SRR577207). Paired-end libraries were filtered for high quality reads of uniform length (100 nt), yielding 25 million read pairs from the 175 and 475 bp libraries, and 15 million pairs from the 750 bp insert library. Paired-end genomic reads were aligned (mapped) to the lettuce chloroplast reference sequence (GenBank accession DQ383816.1) using the CLC Genomics Workbench with stringent parameters (Mismatch cost = 2; Insertion cost = 3; Deletion cost = 3; Length fraction = 0.9; Similarity fraction = 0.9) to find sequences that mapped to the chloroplast genome. Unmapped read pairs were collected and compiled into separate files for downstream mitochondrial assembly and analysis. Reads from mate-pair libraries were used without filtering against the lettuce chloroplast genome.

*L. serriola* acc. US96UC23, whole genome sequence (WGS) paired-end and mate pair genomic libraries were prepared using the DNeasy Plant Mini Kit (Qiagen) from DNA isolated from dark grown seedlings. A paired-end 170 bp insert library was constructed utilizing the NEXTflex PCR-Free approach (BIOO Scientific). The 2.5 kb library was constructed using the Mate Pair Library v2 Kit (Illumina). The 10 kb library was constructed using the Nextera Mate Pair Sample Preparation Kit (Illumina). The raw reads were submitted to the SRA database under accession numbers SRX5097892 (170 bp), SRX5097891 (2.5 kb), and SRX5097890 (10 kb).

Illumina paired-end read data for **L. saligna** (insert size 500 bp) were generated as a part of the International Lettuce Genomics Consortium (ILGC) project (http://lgr.genomecenter.ucdavis.edu/). Mitochondrial reads used in this analysis are available at NCBI Sequence Read Archive under accession number SRX5131542.

### Dovetail Hi-C libraries

Hi-C libraries from leaves were generated by Dovetail™ using their standard proprietary protocol. Libraries were sequenced using an Illumina HiSeq 4000 at the UC Davis Genome Center. Paired reads were deposited in NCBI GenBank SRA under accession number SRX3973834.

### Preliminary Illumina assembly of the mitochondrial genome for **L. sativa**

The plant mitochondrial RefSeq sequences from GenBank were used as a reference to mine mitochondrial contigs from a preliminary assembly of **L. sativa** Illumina reads that was generated using Velvet (89). This preliminary assembly involved six rounds of independent assemblies using Velvet with k-mer values 51, 57, 63, 67, 71, and 75. The Velvet contigs were then joined using CAP3 (90) specifying -o 200 -p 95. A self-BLAST was performed on the CAP3 contigs and a non-redundant set of sequences was selected for downstream analysis.

### Optimization of CLC assembly parameters with **L. sativa** chloroplast PacBio genomic reads

The reference chloroplast genome (GenBank accession DQ383816.1) was used to recover all PacBio reads that contained chloroplast fragments for input into the assembly with the CLC Genomics Workbench using the default approach of the Genome Finishing Module. Assembly parameters were adjusted until we achieved the expected three contigs for this genome, corresponding to the long and short unique segments and the inverted repeat.

### Assembly of the **L. sativa** mitochondrial genome with PacBio reads

The preliminary Illumina-based mitochondrial genome assembly for **L. sativa** was used to query and select PacBio reads containing mitochondrial sequences from the raw PacBio genomic read set. This was achieved using a BLAST-N search of PacBio reads against the Illumina assembly and filtering for contiguous alignments with at least 500 nt and 80% identity. The Illumina reads had already been filtered to remove any plastid DNA reads. The total number of PacBio reads selected for the assembly was adjusted to achieve 250x coverage (∼4,000 reads) before input into the first round of assembly with the CLC Genomics Workbench using the approach and parameters identified from the chloroplast genome assembly. After the first round of assembly, chloroplast segments contained in the mitochondrial assembly were masked and the masked contigs were used to mine additional PacBio reads for mitochondrial sequences for a second round of assembly. This process was repeated until all reads containing mitochondrial segments had been recovered.

### Assembly of the **L. saligna** mitochondrial genome with PacBio reads

The **L. sativa** PacBio mitochondrial genome assembly was used to select PacBio reads from the **L. saligna** data that contained mitochondrial fragments. As before, the amount of reads was adjusted to ∼250x coverage (∼6,000 reads) for input into the first round of assembly with the CLC Genomics Workbench, following the same approach as used for **L. sativa**. The PacBio reads were slightly shorter for **L. saligna**, requiring more total reads to achieve ∼250X coverage.

### Contig naming convention

Upon assembly and preliminary comparison of contigs between species, the following convention was chosen to assign IDs to simplify downstream data analysis and interpretation. Contig naming started with K to avoid the first letters of the alphabet that could be used to designate figure panels on data presentation. Letters X and Y were skipped because of confusion with human chromosomes and letter O because of visual similarity with 0 (zero). **L. sativa** contigs M and N are frequently adjacent on genome isoforms, so alphabetical ordering keeps them together in data tables. Two letters were assigned to the longest **L. saligna** contig UV because it can be decomposed into two distinct sequences that are similar to **L. sativa** U and W. At the same time, **L. sativa** W is different from **L. saligna** V by having extra segments and thus was “wider.” Alphabetical sorting keeps them (U + V) in the right order. For one repeat, the letter R was assigned, and for the shortest one T (tiny). **L. saligna** S contig stands for “special” (integrated linear plasmid). These simple mnemonic rules were helpful for navigating the diverse data in the process of genome assembly because so many steps had to be visualized and resolved manually. This letter coding style was particularly useful in analysis of data using MS Excel spreadsheets and conditional formatting that allowed distinct coloring for different contigs and their segments.

### Inference of mitochondrial secondary building blocks using raw PacBio reads

Contiguity of the assemblies (sequential order of contigs or primary structural units) was determined based on the analysis of full length raw PacBio reads. This led to the set of secondary building blocks that were used to infer the comprehensive set of mitochondrial genome isoforms.

For the reverse PacBio read mapping approach, a sliding window of 2 kb with a step size of 1 kb was applied to each contig of the assembly (S9A Fig) to generate a library of overlapping fragments. Overlapping tiling fragments were mapped/aligned back to raw mitochondrial PacBio reads. A BLAST-N search was used to map tiling fragments to PacBio reads using the relaxed parameters to allow longer gaps (compared to default parameters) over contiguous alignment (-V T -F F -e 1e-60 -y 50 -X 75 -Z 500). A custom BLAST-N parser (https://github.com/alex-kozik/atgc-tools/blob/wiki/tcl_blast_parser.md) was used to generate a tab-delimited table that was used for sequential order analysis of primary structural units in PacBio reads. PacBio reads containing at least 10 distinct tiling fragments of the mitochondrial non-redundant contiguous genome sequence for **L. sativa** and 12 distinct tiling fragments of the mitochondrial non-redundant contiguous genome sequence for **L. saligna** were selected and compiled into maximally informative sets. Maximally informative sets resulted in 4,046 and 11,009 PacBio reads for **L. sativa** and **L. saligna**, respectively. These sets were used for the analysis of junctions between different mitochondrial genome primary structural units. The distributions of junctions between different units were first analyzed visually with MS Excel (S9B Fig). Regular expressions with egrep were utilized to quantify the junctions from tables in tab-delimited format.

### Re-assembly and validation of junctions

Regions of PacBio reads specific to individual junctions (as identified in each set of secondary building blocks, see above) were compiled as separate, small subsets and reassembled to recover the precise sequence at each junction. For each junction, PacBio reads containing termini of two contigs were trimmed to 4 – 6 kb so the junction region was located in the middle of the trimmed reads. Subsets of the trimmed PacBio reads were assembled with CLC for each junction individually. The alignment of mitochondrial contigs to reassembled junction regions allowed for the determination of precise sequences between mitochondrial contigs.

### Error correction using Illumina reads

Paired-end Illumina reads for **L. sativa** and **L. saligna** were used for error correction of the assembled PacBio contigs. Illumina reads from genomic libraries were mapped/aligned to the PacBio contigs to correct homopolymer errors and consensus sequences were selected based on what was supported by the majority of the Illumina reads.

### Mitochondrial contig stoichiometry

The stoichiometry of mitochondrial contigs was determined by PacBio read coverage. A sliding window of 2 kb with a step size of 0.5 kb was applied to each contig of the assembly to generate a library of overlapping fragments. BLASTN was performed on each 2 kb tiling fragment versus the set of the most informative PacBio reads. To calculate the coverage, the number of PacBio reads for each tiling fragment was counted if the alignments were longer than 1.8 kb with an identity of 80% or better. Since all non-tandem repeats of medium size are shorter than 1.2 kb, alignments due to repeated sequences were not included. The total number of alignments per tiling fragment were plotted (this reflects coverage per segment across contig) and compared to each other.

The coverage for each junction was calculated for the **L. sativa** and **L. saligna** mitochondrial genomes following methods similar to those above. The junction-spanning assemblies, consisting of 2 kb fragments with 1 kb of sequence on either side of the junction, were used as BLAST queries versus the PacBio reads to verify the assembly and provide numerical values for the fractionation of each junction in the pool of PacBio reads. Each segment that corresponded to a particular junction generated one of two types of alignment. The first type was a completely uninterrupted alignment that was specific for a particular junction. The second type was a fragmented alignment that corresponded to junction components in different locations of the mitochondrial genome. The alignments of different lengths were plotted for each BLAST hit (sorted by alignment length). The transition from complete (1.8 kb) to shorter segmented alignments indicated the fraction of each junction type in the pool of PacBio reads.

### Isoform inference and quantification

Mitochondrial contigs generated with the CLC assembler were considered as primary units after polishing. Junctions between primary structural units inferred by reverse PacBio read mapping and analysis (see above) determined their sequential order. Visual inspection of secondary building blocks and analysis of their possible inter-connections taking into account their stoichiometries resulted in the set of mitochondrial genome isoforms.

In this paper, we define contigs as primary structural units; secondary building blocks are combinations of several primary structural units (as detected by reverse read mapping); isoforms are combinations of several secondary building blocks with stoichiometry taken into account. Stoichiometry data were derived from contig coverage with PacBio and Illumina reads.

### Validation of the assemblies with Illumina mate-pair libraries

For **L. sativa** and *L. serriola*, we used Illumina mate-pair libraries to validate structural accuracy of the PacBio assembly and inferred mitochondrial isoforms. Illumina mate-pair reads from genomic libraries of **L. sativa** with insert sizes of 2.5 and 10 kb were aligned to the assembled isoforms using BWA (91). The distances between aligned mate-pair reads were calculated based on their positions/coordinates on the mitochondrial reference sequence. Pairs with distances greater than 1 kb were selected and compiled into subsets of equal size (320,000 pairs) for each library. Two dimensional distance heat plots for 1 kb bins were generated using modified Python scripts (https://github.com/alex-kozik/atgc-uni-cluster) and the Pixelirator visualization program (http://cgpdb.ucdavis.edu/data_pixelirator/).

### Higher order arrangement analysis with Hi-C libraries

Read pairs from Hi-C libraries were filtered to eliminate ligation artifacts that do not reflect true long-distance interactions. The use of the *Mbo*-I restriction enzyme for fragmentation prior to ligation creates GATC-GATC dimer sites upon ligation of the *Mbo*-I digested termini, which do not exist in the original mitochondrial genome. Reads with GATC-GATC sites were selected and used for the long distance interaction analysis. Each read with a GATC-GATC site was split into pairs of sub-reads and each read of a split pair was mapped separately to the mitochondrial reference. The distances were calculated as in the mate read analysis. A subset of 640,000 pairs with distances greater than 1 kb was randomly selected for the distance analysis and visualization.

### Repeat analysis

Repeated sequences were found as described previously (25) using BLAST-N with a word size of 50, ungapped, no masking, reward +1, penalty -20, and e-value 1,000. These parameters identified nearly identical repeats, indicating either a recent duplication, or recent homologous recombination and gene conversion of the two copies. Less similar repeats were presumed to have mutated and drifted without recent gene conversion, indicating that they are no longer engaging in productive homology-based events.

### Detection of rare recombination events between repeats less than 1 kb

For the detection of rare recombination events between short repeats, all available mate pair reads of insert size 5 and 10 kb for **L. sativa** and 10 kb for *L. serriola* were used. The mitochondrial fraction of these libraries had ∼1 million pairs per library for **L. sativa** and ∼0.7 million for *L. serriola*. Mate-pair reads were aligned to the genomic sequences using BWA (91), the same parameters as above, and masking the second copy of the large repeats in the reference genome. Two-dimensional distance heat plots for 1 kb bins were generated as described above. Recombination events between short repeats resulted in distinct patterns of double short diagonals that are separated from the main diagonal. Variable intensity of the short diagonals was an indication of recombination frequency for particular repeats. Numerical values were analyzed in an MS Excel table as shown in S5D Fig.

### Gene content, analysis, and mitochondrial genome annotation

The mitochondrial genome was annotated for expected mitochondrial features using the Mitofy web server (http://dogma.ccbb.utexas.edu/mitofy/). In order to find coding sequences missed by Mitofy, all open reading frames were manually examined. The two different junction open reading frames for *atp1* and *rps4* were translated and compared to other plant mitochondrial atp1 and rps4 protein sequences using BLAST-P (https://blast.ncbi.nlm.nih.gov/Blast.cgi). Data files for the GenBank submission were compiled using GB2sequin (92).

### Sequence analysis with MUMmer

Mitochondrial genome sequences or their segments were compared with each other using MUMmer4 (93). Alignments in MUMmer delta format were generated with nucmer utility with options ‘--maxmatch --nosimplify.’ SVG plots were created with the mummerplot program and custom edits of corresponding input files for gnuplot (http://www.gnuplot.info/).

### Leucaena trichandra dataset

The *L. trichandra* dataset (74) was used to validate our approach to mitochondrial genome assembly with a different species. *L. trichandra* mitochondrial PacBio reads (SRX2719625), Illumina 4 kb mate-pair libraries (SRX2719623 and SRX2719624), and the putative autonomous mitochondrial DNA element (MH717174.1) were reanalyzed as described above and compared to the published *L. trichandra* mitochondrial genome assembly (MH717173.1).

Isolation of mitochondria and preparation for in-gel fluorescence microscopy and pulsed-field gel electrophoresis Whole seven-day-old seedlings (roots and shoots) of **L. sativa** cv. Salinas grown in the dark under sterile conditions at 15°C were harvested and 5 – 10 g of plant material was flash frozen in liquid nitrogen. The frozen tissue was ground to a fine powder using a mortar and pestle, then ground again after adding 5 mL of High Salt Buffer (HSB: 1.25 M NaCl, 40 mM HEPES pH 7.6, 2 mM EDTA pH 8, 0.1% Bovine Serum Albumin (BSA), 0.1% 2-mercaptoethanol) until the frozen slurry melted. The liquid homogenate was filtered through 1 layer of Miracloth, then centrifuged at 4°C at 3,000 x g to pellet chloroplasts and nuclei. The supernatant was transferred to a fresh set of tubes and centrifuged at 4°C at 20,000 x g to pellet mitochondria. The mitochondrial pellet was resuspended in Liverwort Dilution Buffer (LDB; 0.4 M sorbitol, 1 mM EDTA, 0.1% BSA, 20 mM HEPES-KOH, pH 7.5) at half the volume of HSB used for the initial grinding and centrifuged at 4°C at 3,000 x g to again remove chloroplasts and nuclei by pelleting. The supernatant was transferred to a fresh set of tubes and centrifuged at 4°C at 20,000 x g to pellet mitochondria. The mitochondrial pellet was resuspended in half the original volume of LDB and the 3,000 x g centrifugation, transfer of supernatant and 20,000 x g centrifugation were repeated. The mitochondrial pellet was resuspended using 2 μL of LDB per μL of pelleted mitochondria. For PFGE, low melt point agarose (LMPA) and sorbitol were added to the suspension to achieve final concentrations of 0.7% LMPA and 0.41 M sorbitol before pipetting into an agarose plug mold, which was allowed to cool for 20 min at 4°C. For in-gel fluorescence microscopy, agarose plugs of the mitochondria were prepared as for PFGE, but the mitochondria were first diluted 1:20, 1:100, or 1:200 in LDB before embedding. Samples for PFGE were run on a 1.5% agarose gel in 0.5X Tris-Borate-EDTA (TBE) buffer at 5 V/cm with a pulse time of 30 s for 24 hours and stained in 3X Gel Red (Biotium) before imaging. For in-gel fluorescence microscopy, ¼ of an agarose plug was soaked in BET solution (3% 2-mercaptoethanol, 0.1 μg/mL EtBr, 1X TBE) for 30 mins. The BET solution was removed and the sample was soaked in fresh BET before slide preparation. Alternatively, ¼ of an agarose plug was soaked in GET solution (3% 2-mercaptoethanol, 0.5x QuantiFluor® dye (Promega), 1X TBE) for 30 mins. To prepare slides, ½ of the stained agarose plug was placed on top of a glass slide on a heat block at 60 – 65°C and mixed with 15 μL ABET (1% LMPA in BET) and a coverslip was placed on top. When the agarose was completely melted underneath the coverslip, the slides were removed from the heat block and sealed with nail polish before microscopy. To measure the circular molecules, the images were analyzed using Fiji software (https://imagej.net/Fiji). The path along each entire circular molecule was measured three times and we used the average of these three measurements.

## Acknowledgments

We thank Luca Comai, Arnold Bendich, and Jeff Palmer for helpful comments on the manuscript and Elizabeth Georgian for editorial assistance. We are also grateful to Pauline Sanders for supplying and maintaining seed stocks and Huaqin Xu for assistance with data submission to NCBI GenBank.

---

**S1 Fig.**
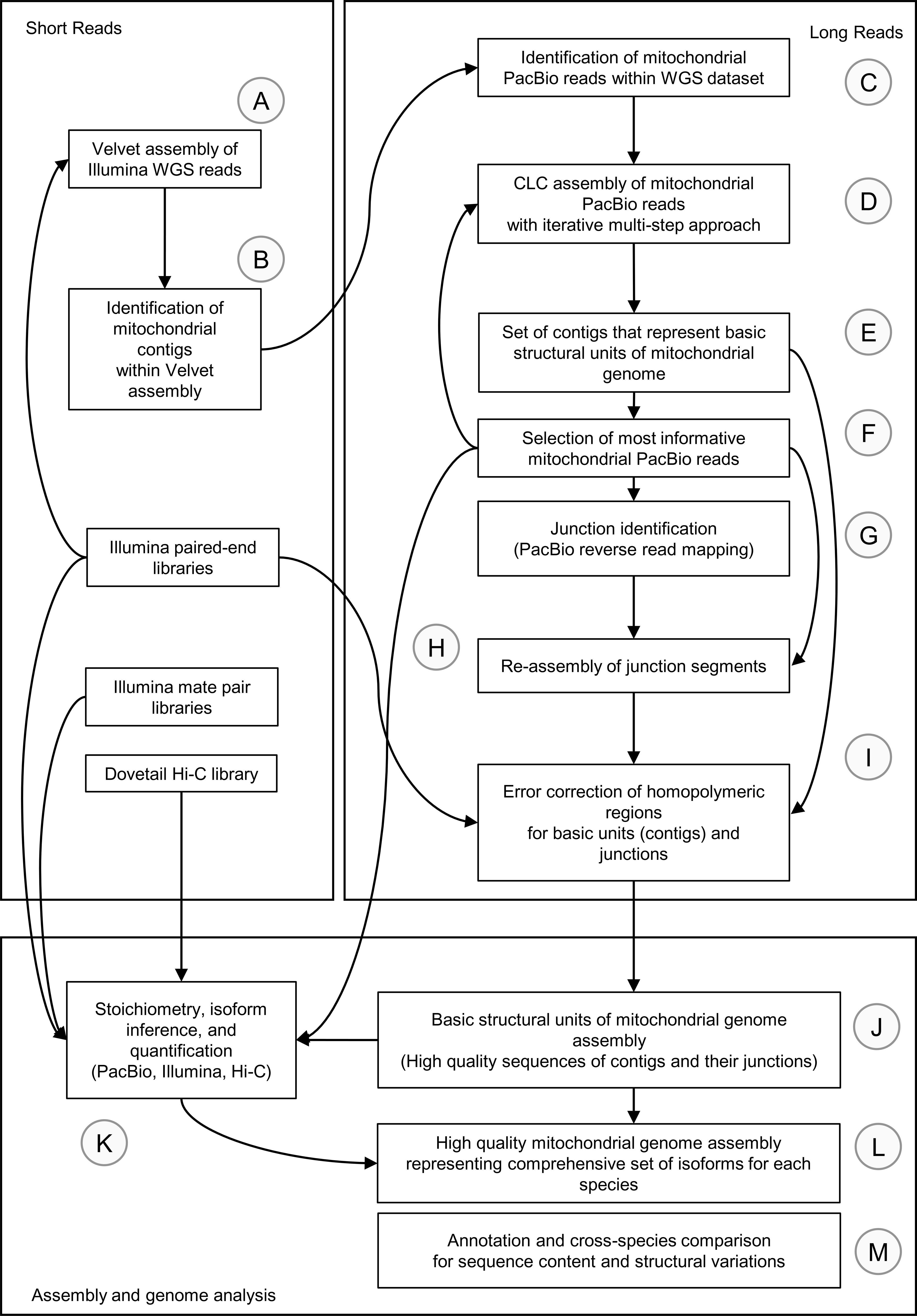
Mitochondrial genome assembly and analysis strategy. The flowchart summarizes all of the key steps of the *Lactuca* mitochondrial genome project along with the source and type of raw data used in each iteration of the assembly and analysis.

**S2 Fig.**
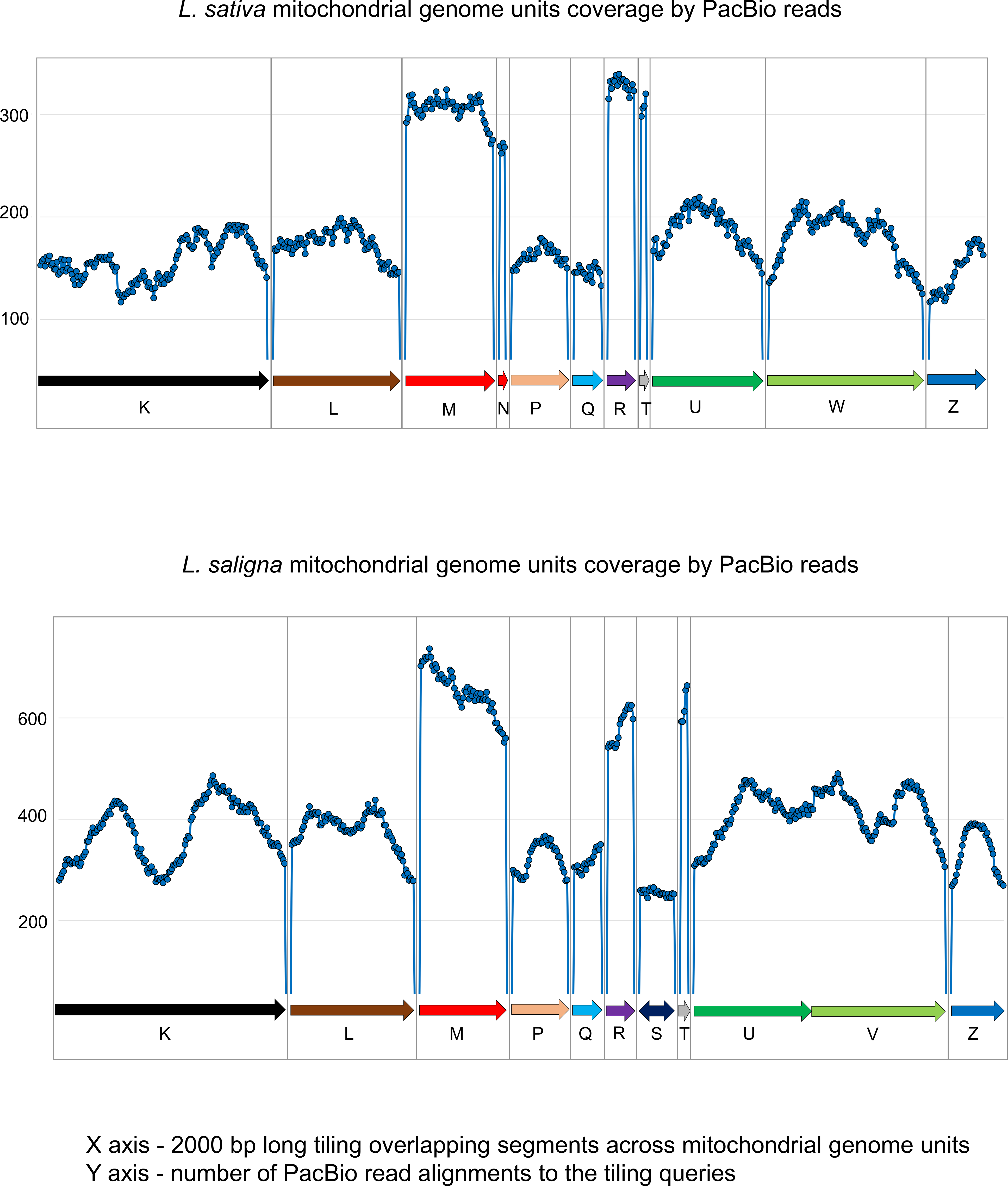
Results of the stoichiometry analysis for primary structural units. A tiling library of overlapping fragments (similar to that used for reverse read mapping) was used to estimate coverage for each fragment (see Contig Stoichiometry in Materials and Methods). The Y axis shows coverage; the X axis displays the coordinates of the primary structural units. Note the elevated coverage in the middle portion in the majority of basic units. Stoichiometry values for each primary structural unit were used for the mitochondrial genome isoform modeling.

**S3 Fig.**
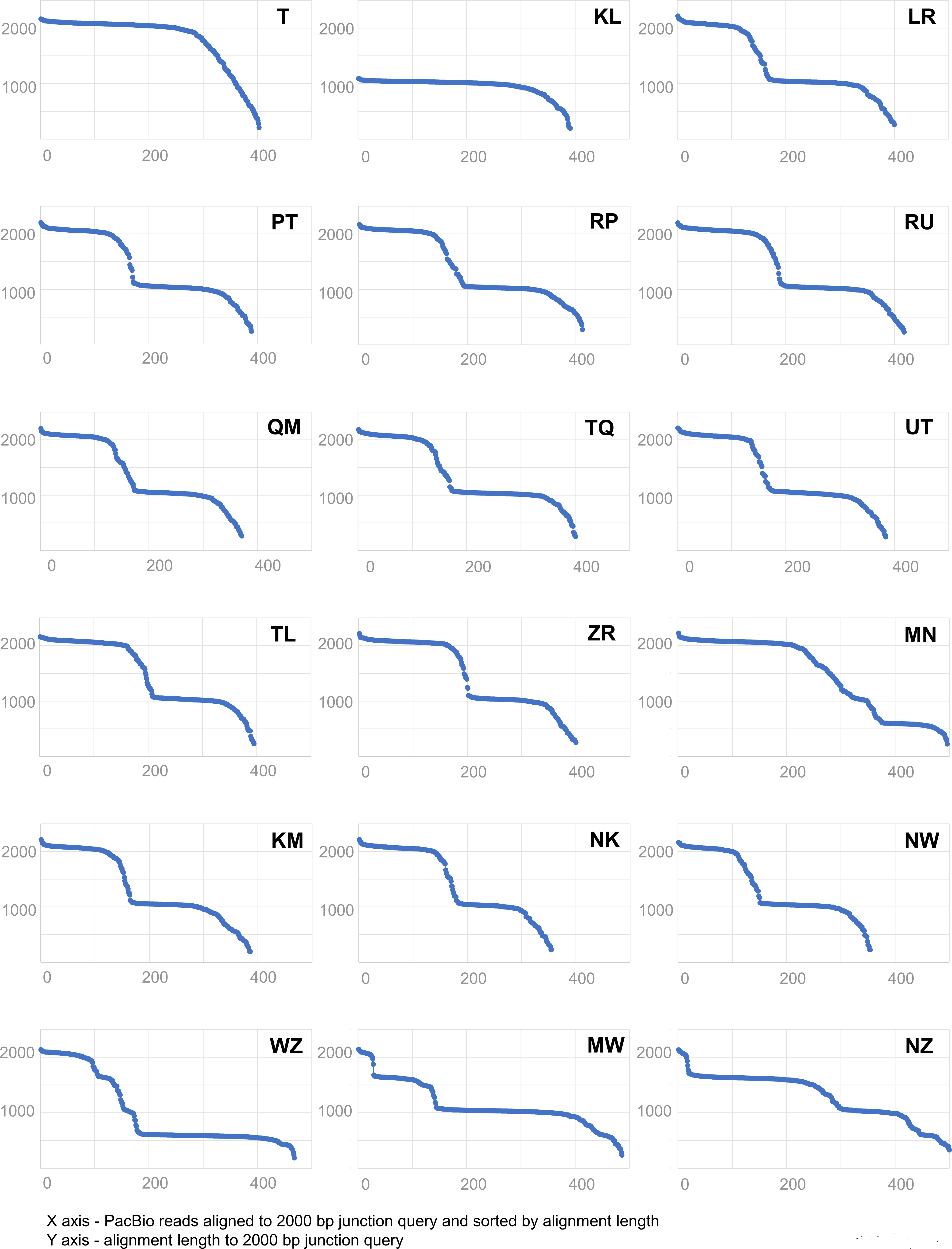
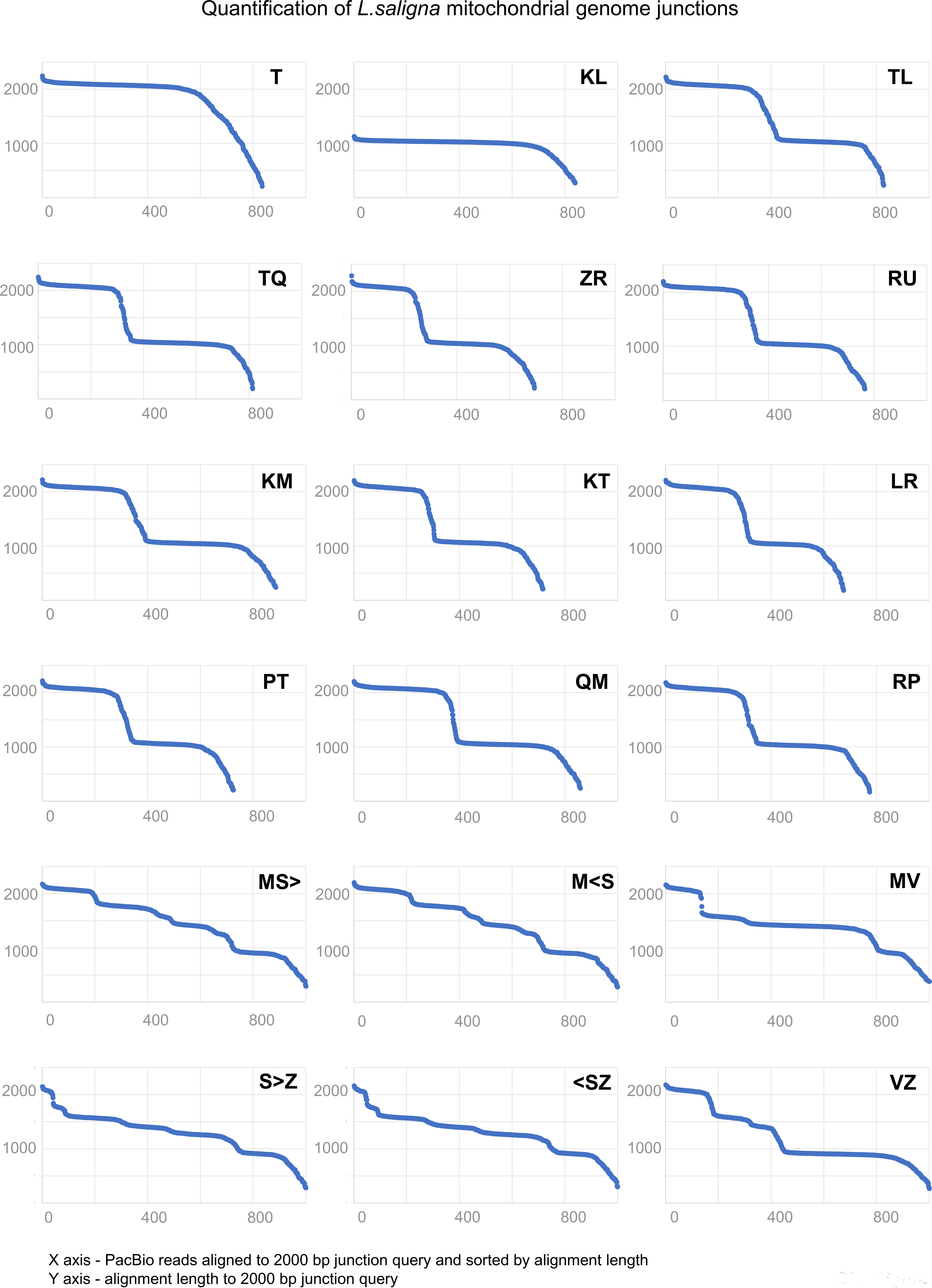
Stoichiometry analysis for each detected junction between different primary structural units. The set (library) of all identified junctions (2 kb each, see Materials and Methods, Mitochondrial Contig Stoichiometry) was analyzed for the fraction of uninterrupted alignments within a pool of most informative PacBio reads for *L. sativa* (A) and *L. saligna* (B). All BLAST-N alignments are plotted on the X axis and sorted according to alignment lengths. The length of each alignment is shown on the Y axis. Breakpoints (transitions between uninterrupted alignments longer than 1.8 kb) are an indication of the abundance of each particular junction in a pool of PacBio reads. This value (number of uninterrupted alignments) reflects the proportion of any particular junction within isoforms of a mitochondrial genome.

**S4 Fig.**
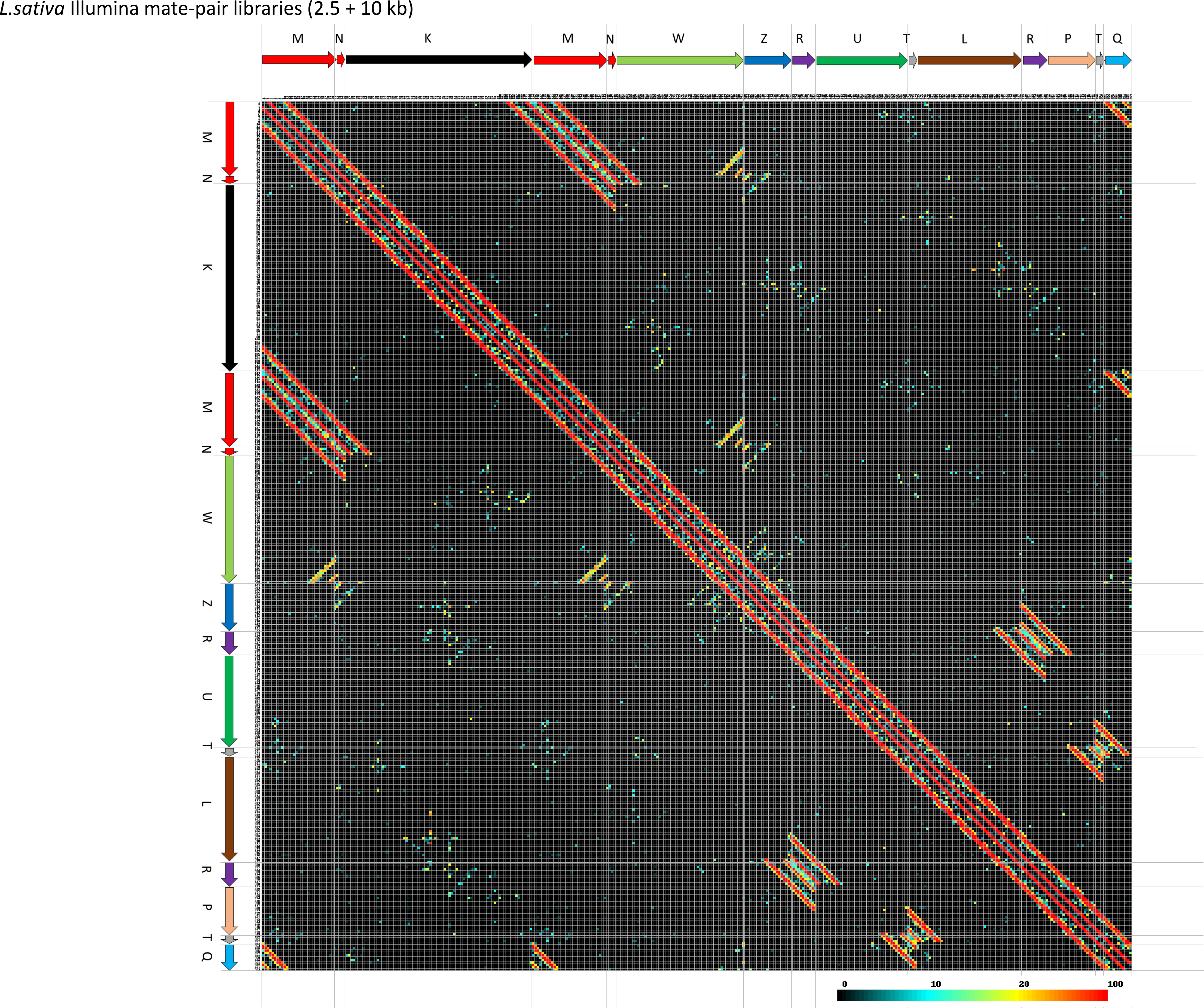
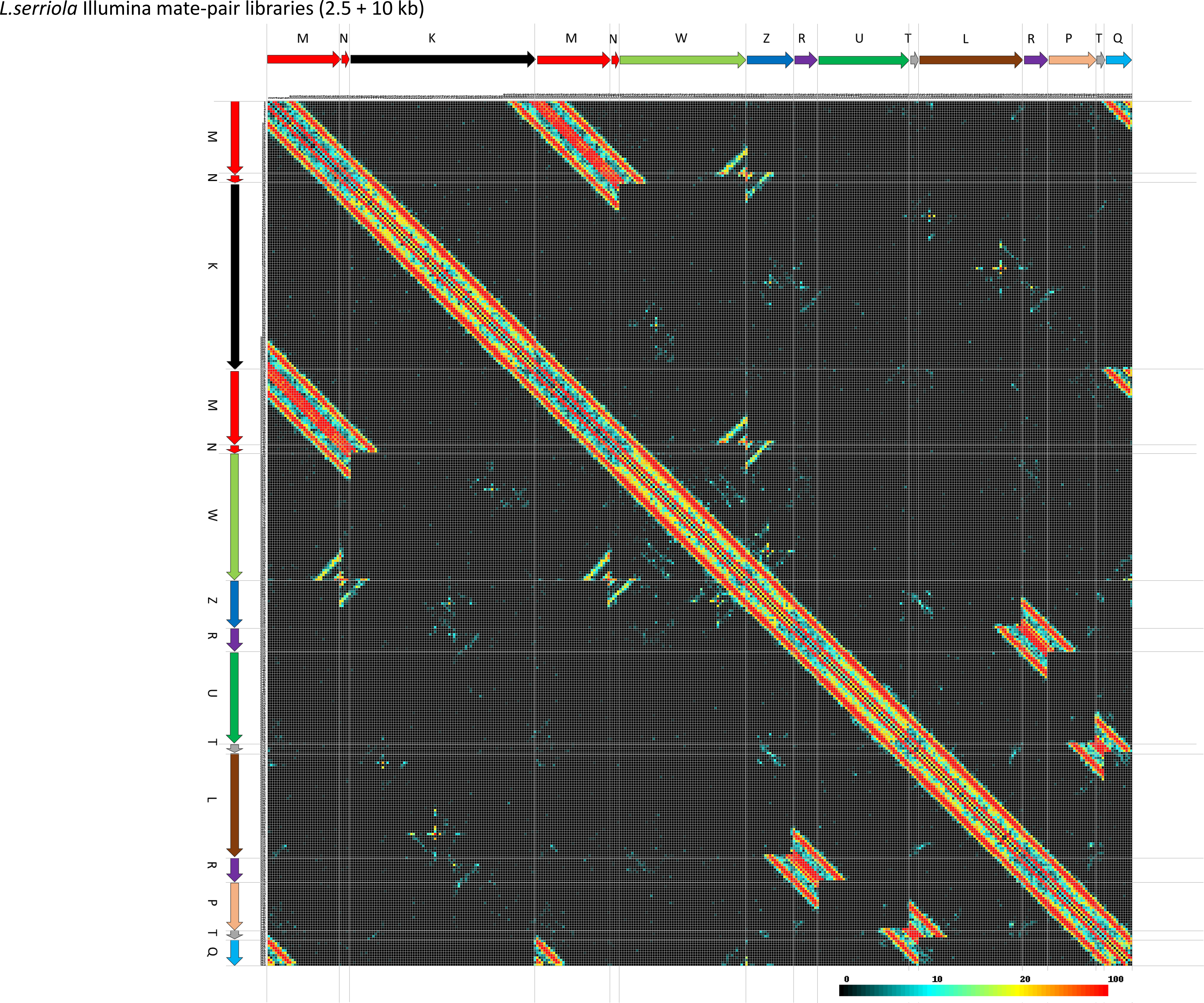
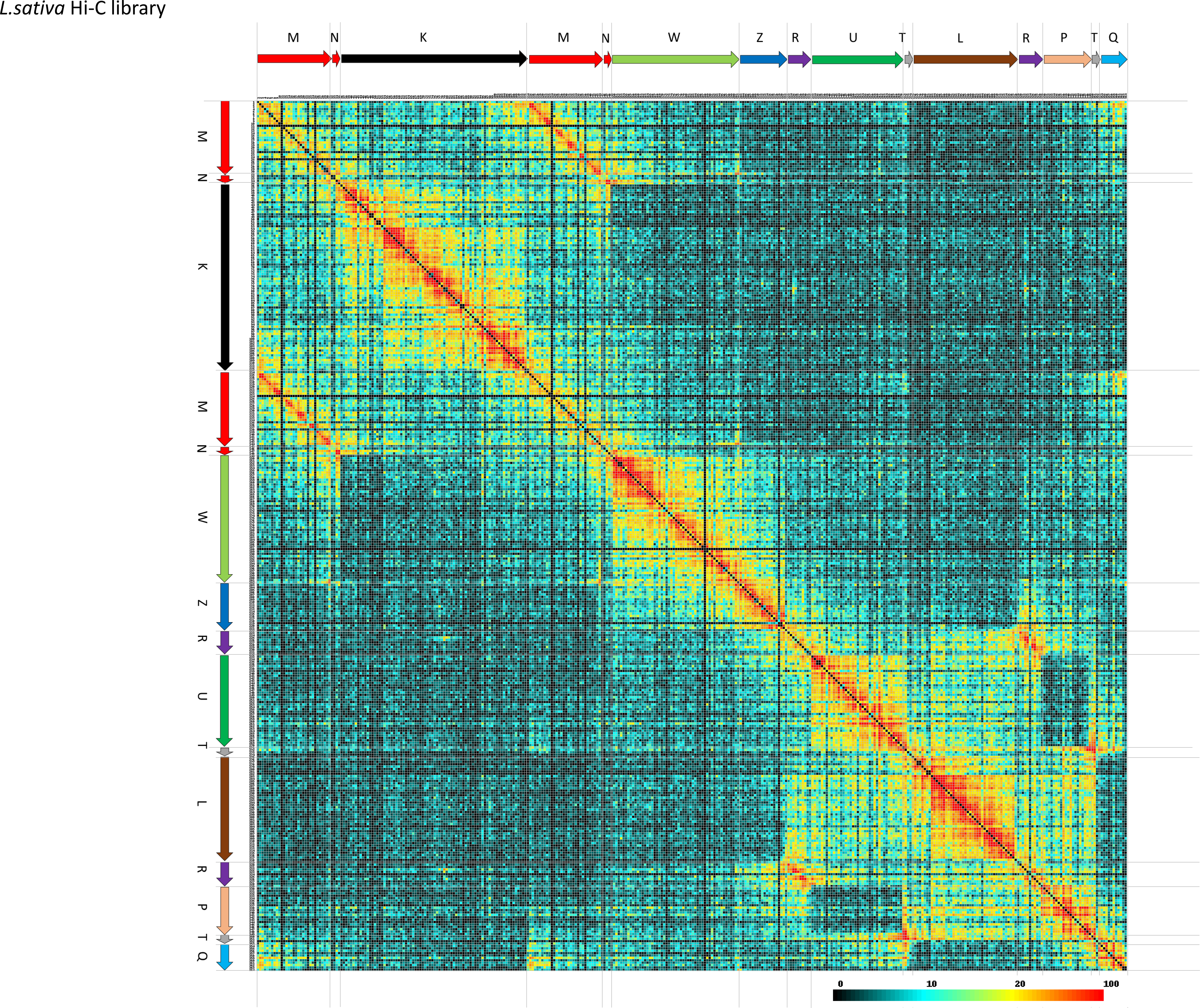
High resolution images shown in Fig 4.

**S5 Fig.**
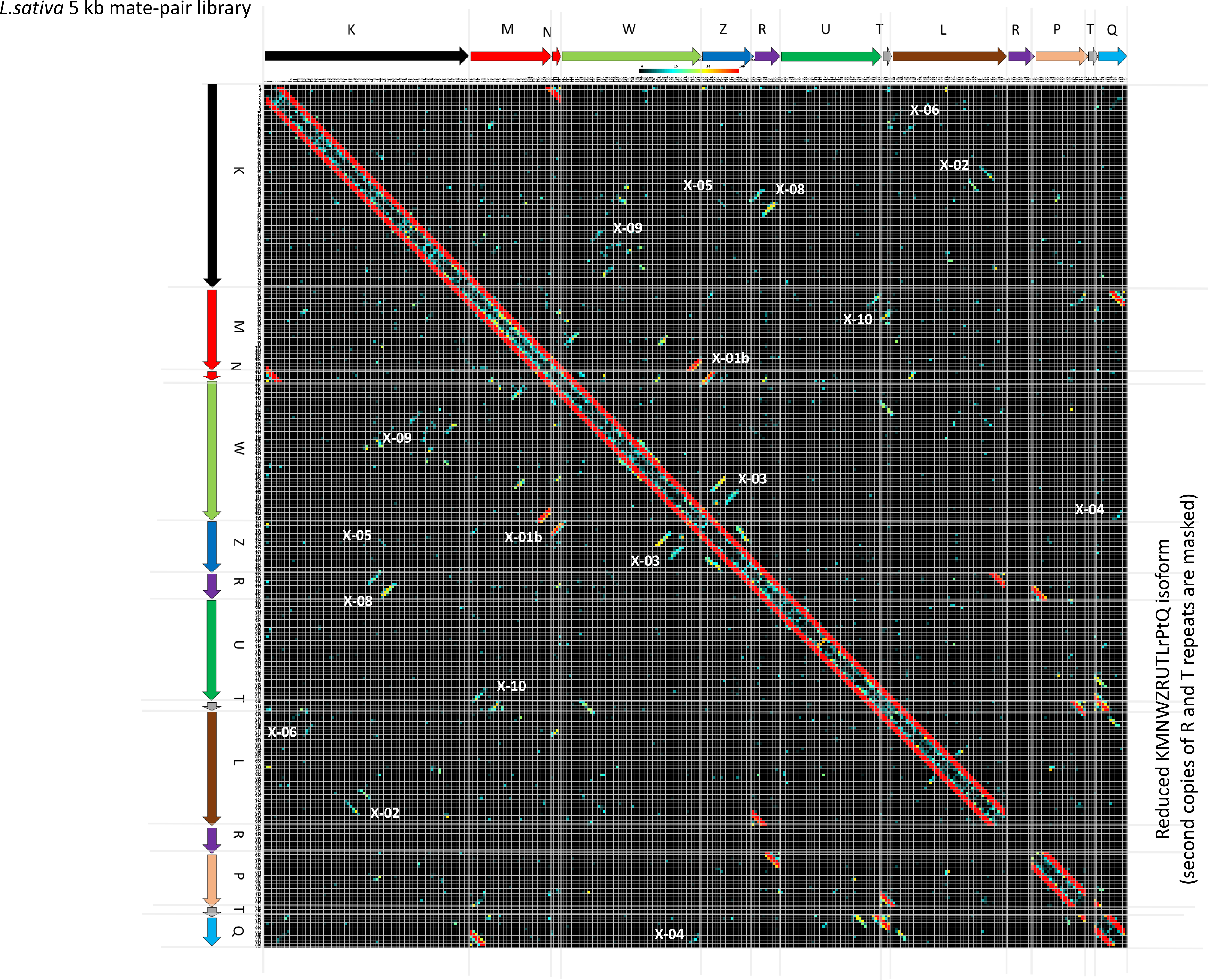
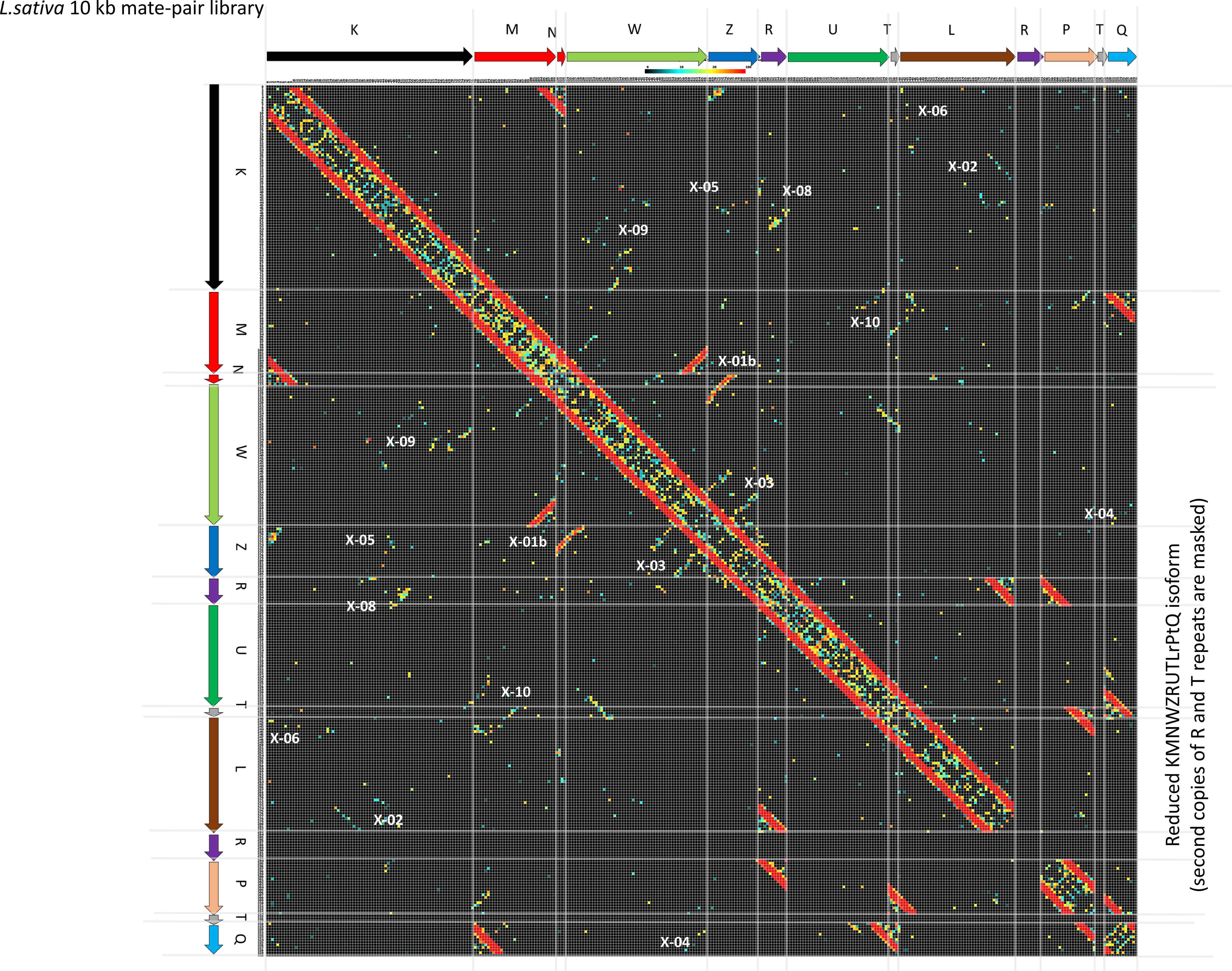
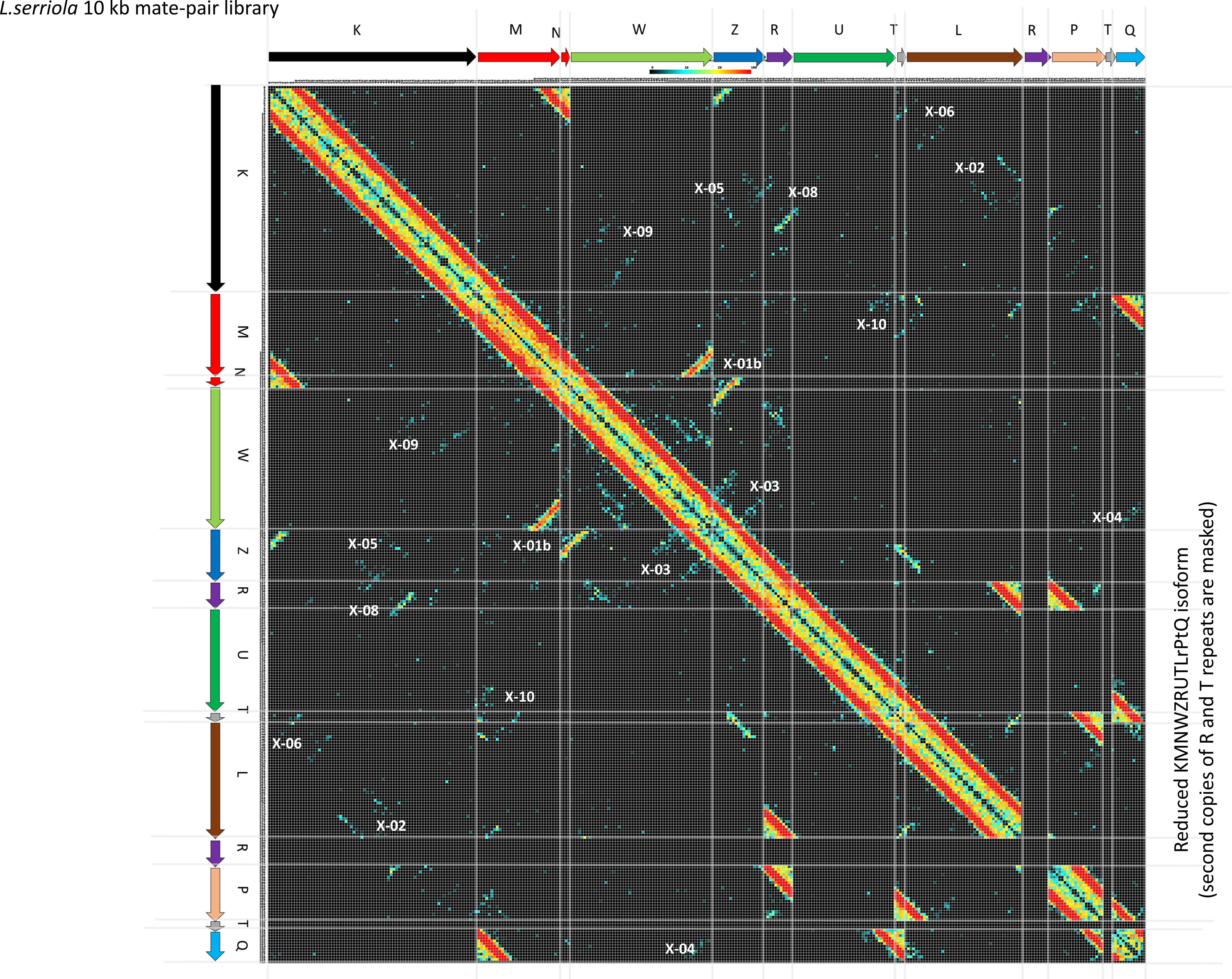
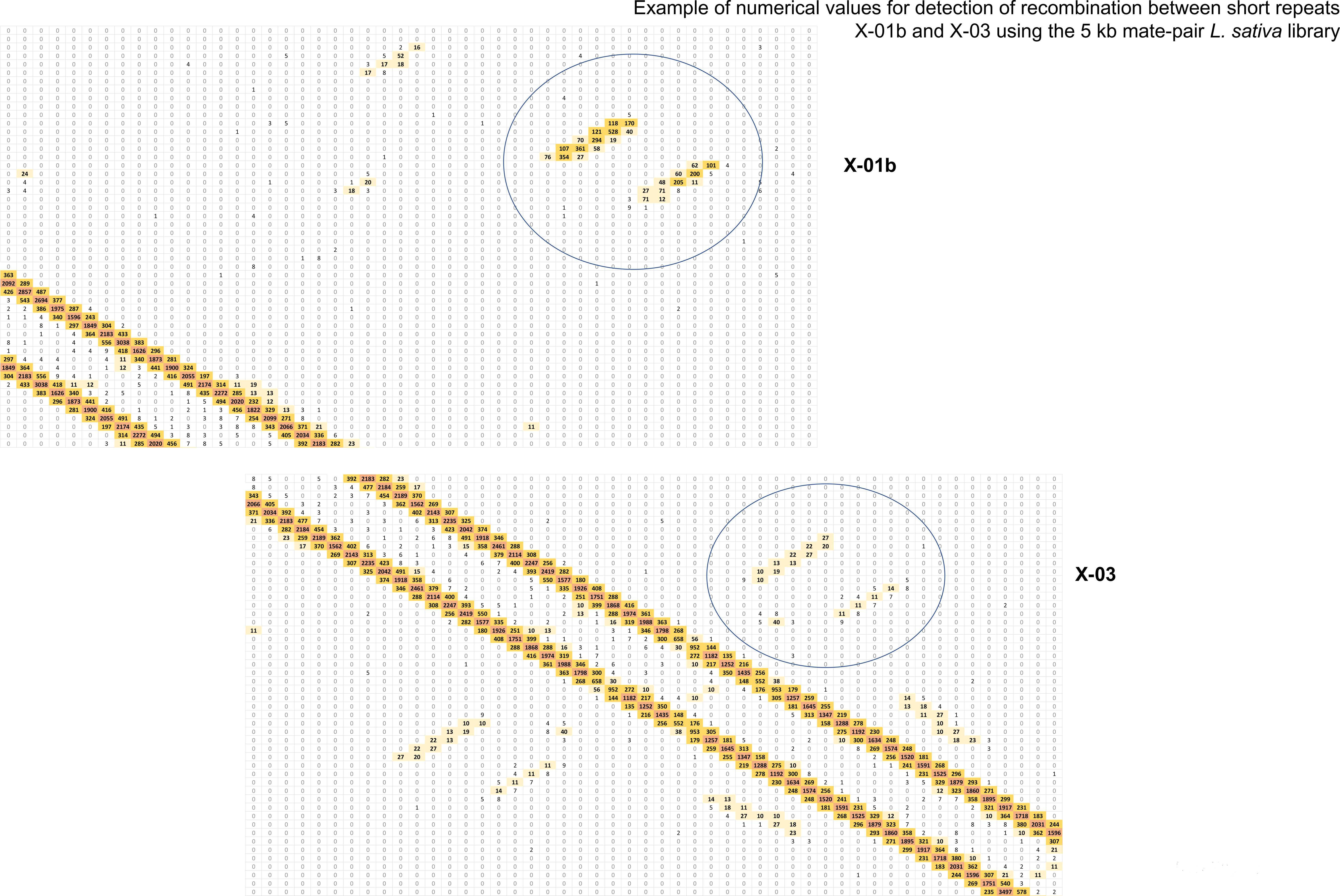
Detection of recombination events at repeats of up to several hundred bp using Illumina mate pair libraries. Plots of distances between read-pairs in 1 kb bins for *L. sativa* mate pair libraries of 5 kb (A) and 10 kb insert sizes (B), and L. serriola 10 kb (C). Only unique read mappings were selected from all available reads for the analysis and data interpretation. S5D Fig: Example of numerical values for detection of recombination between short repeats X-01b and X-03 using the 5 kb mate-pair *L. sativa* library (see S5A Fig for the complete 2D plot). Each cell is a 1 kb bin across the mitochondrial genome isoform α. Values within each cell give the number of times mate-pair reads mapped within the same bin. The majority of mate-pair reads are mapped and equally distributed over the main diagonal and within large repeats (R or T). Rare recombination events between short repeats generate distinct shapes (shown as X-01b and X-03) that are located away from the main diagonal. The values reflect the frequency of recombination. Thus, for repeat X-01b, it could be estimated that corresponding recombination frequency is ∼10% and ∼1% for repeat X-03.

**S6 Fig.**
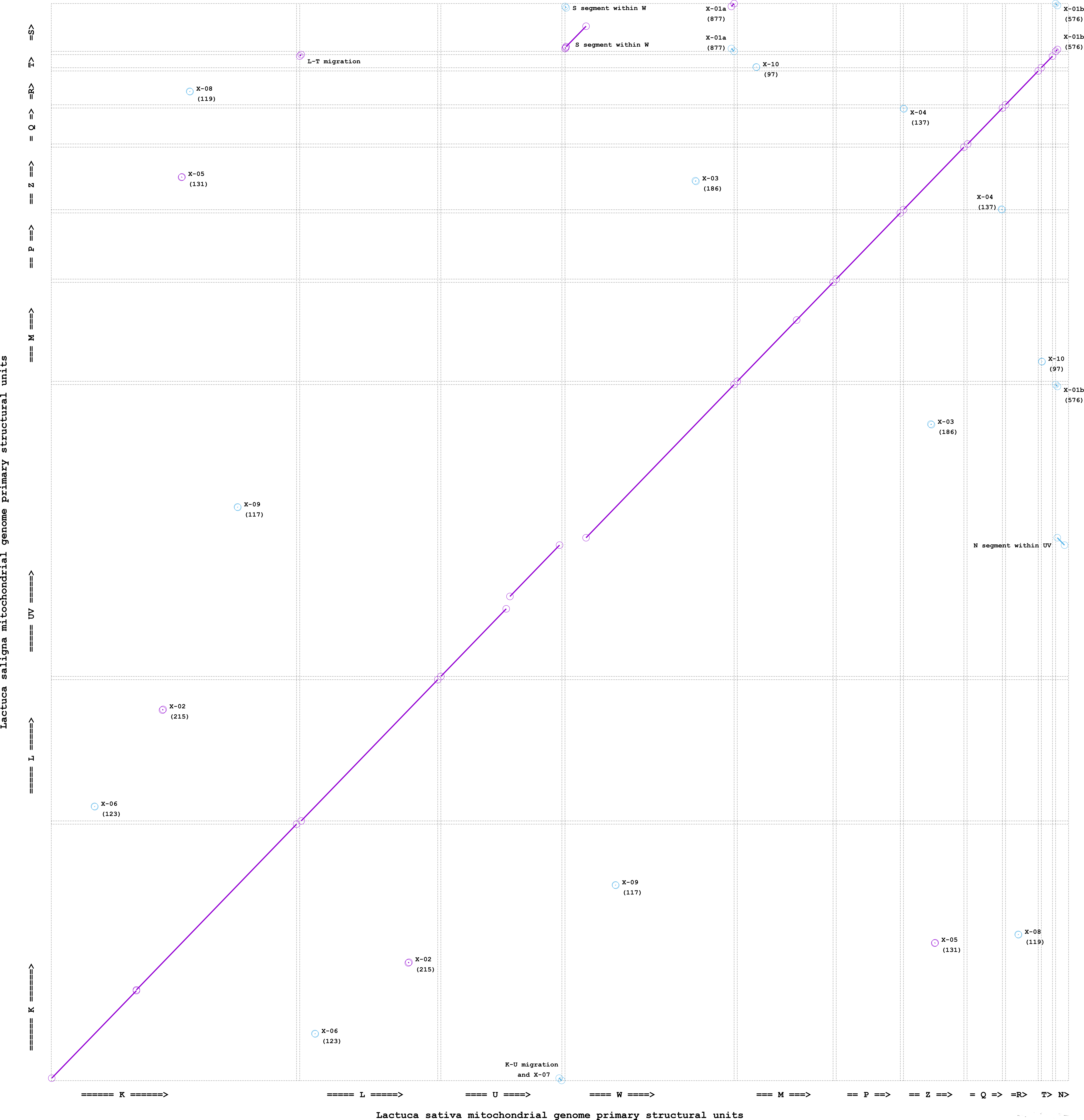
Side by side comparison (MUMmer diagonal plot) of mitochondrial genome primary structural units for *L. sativa* and *L. saligna*. Each unit is represented by an individual contig sequence. Units are separated by 1 kb artificial intervals to simplify visualization of their boundaries. Short repeats are designated by labels with ‘X-’ prefix along with their length in parentheses. Structural rearrangements between basic units (migrated segments) are located outside of the main diagonal and labeled accordingly.

**S7 Fig.**
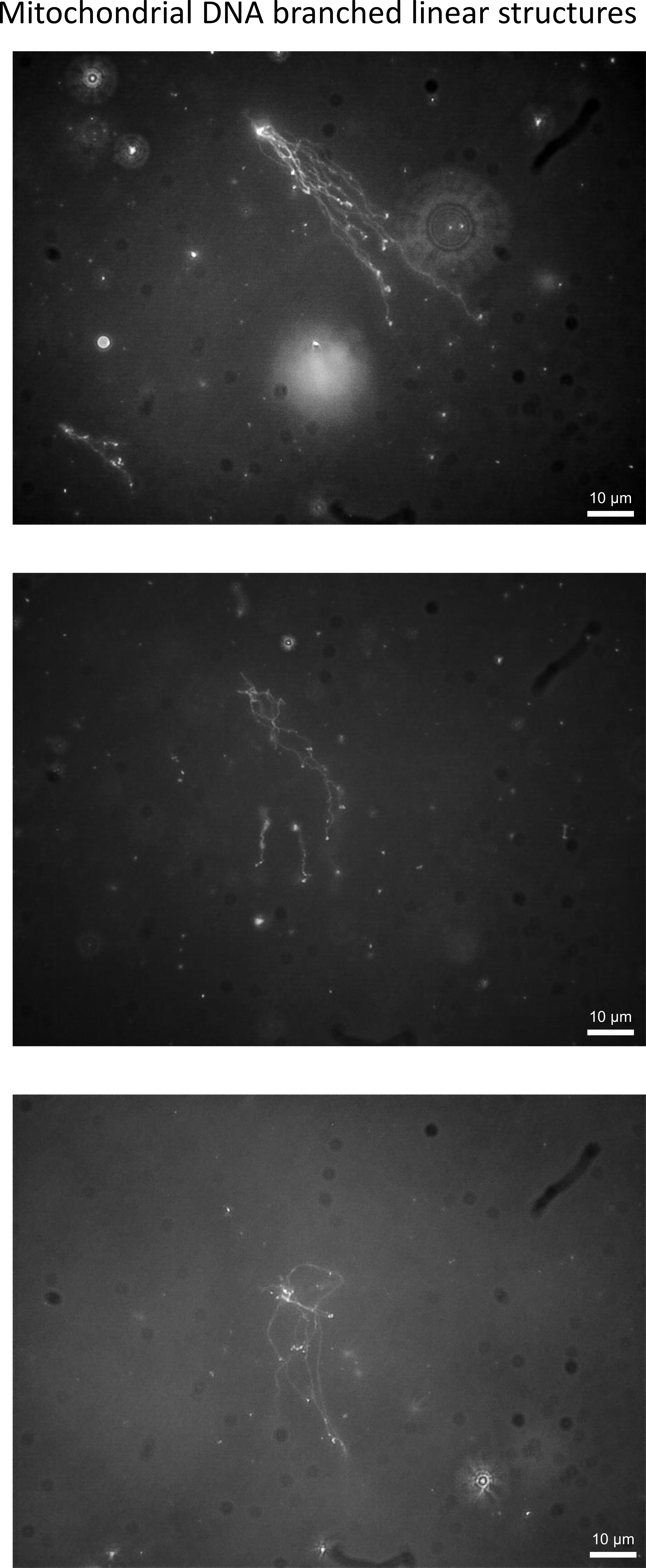
Mitochondrial DNA branched linear structures. Additional examples of molecules scored in Fig 7G.

**S8 Fig.**
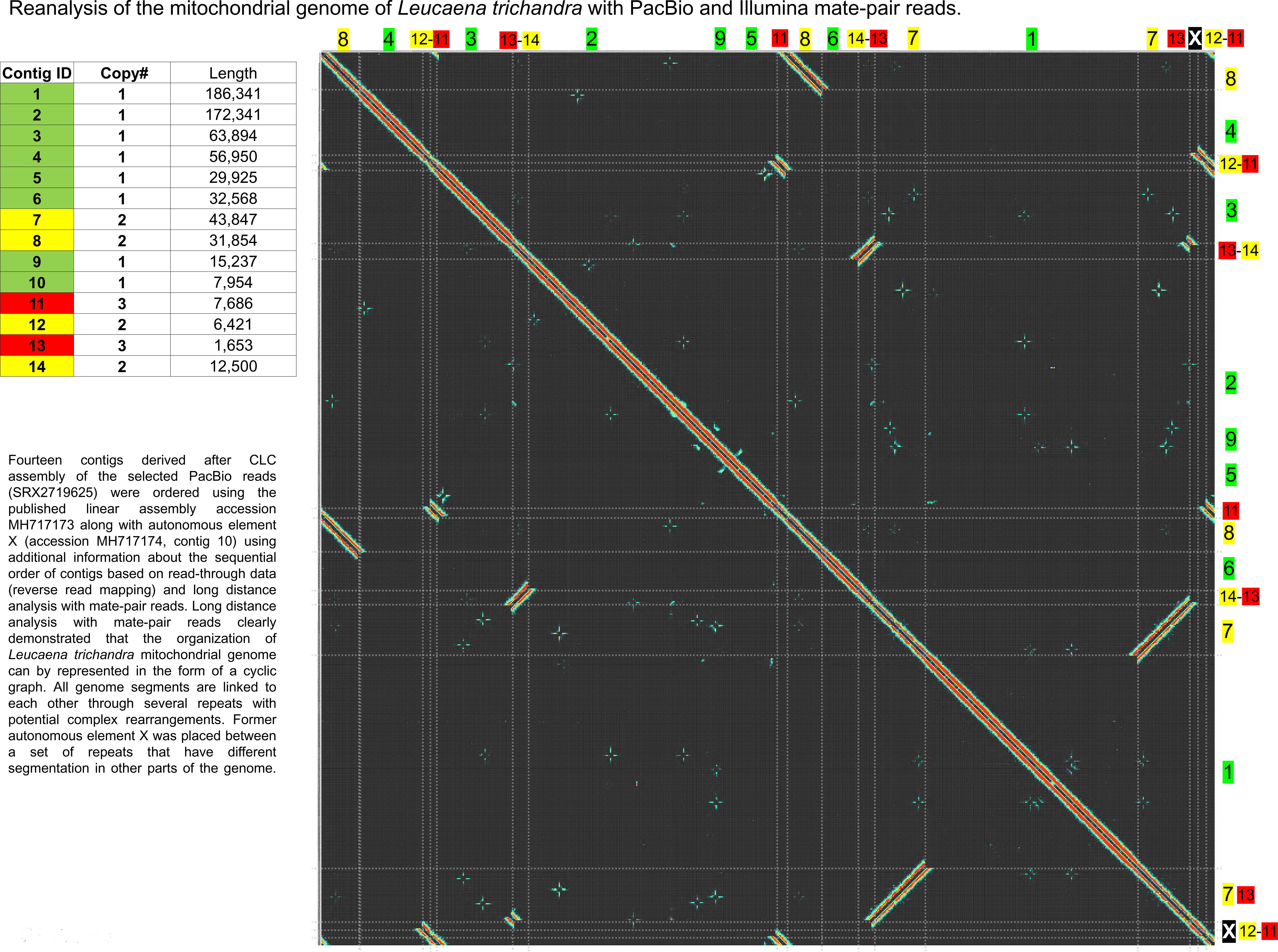
Reanalysis of the mitochondrial genome of Leucaena trichandra with PacBio and Illumina mate-pair reads. Fourteen contigs derived after CLC assembly of the selected PacBio reads (SRX2719625) were ordered using the published linear assembly accession MH717173 along with autonomous element X (accession MH717174, contig 10) using additional information about the sequential order of contigs based on read-through data (reverse read mapping) and long distance analysis with mate-pair reads. Long distance analysis with mate-pair reads clearly demonstrated that the organization of the Leucaena trichandra mitochondrial genome can be represented in the form of a cyclic graph. All genome segments are linked to each other through several repeats with potential complex rearrangements. Former autonomous element X was placed between a set of repeats that have different segmentation in other parts of the genome.

**S9 Fig.**
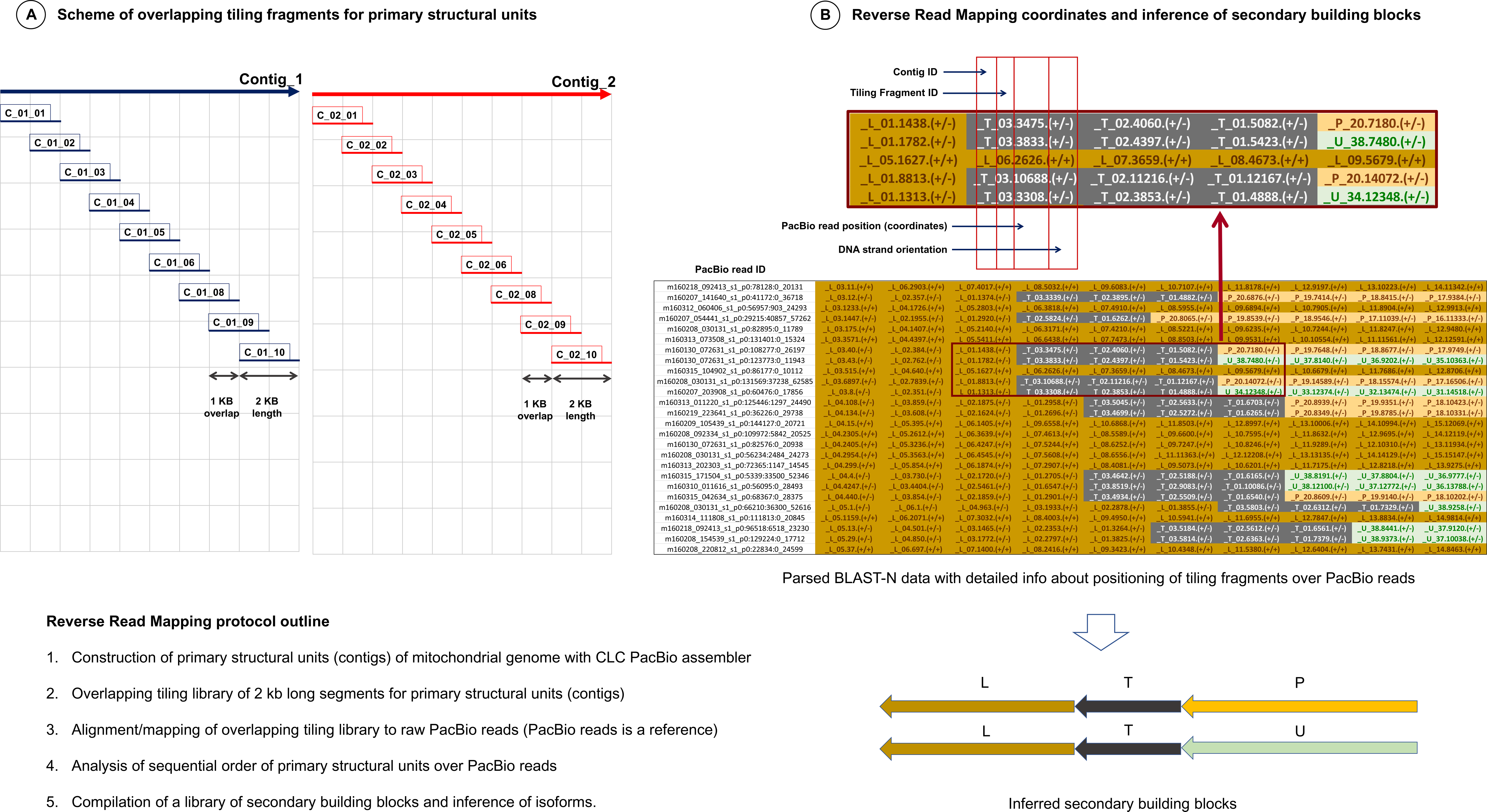
Summary of reverse read mapping approach. Explanation and data interpretation of the reverse read mapping approach. Panel A: Scheme of overlapping tiling fragments for primary structural units and reverse mapping protocol outline. Tiling fragments were used as queries in BLAST-N searches versus a database of mitochondrial PacBio reads. Results of BLAST-N were parsed and exported into an MS Excel table as shown in (B). Visual inspection of the distribution of primary structural units over long PacBio reads in an MS Excel table with subsequent search queries on text files provided information about the sequential order of primary structural units within the PacBio reads. This ultimately led to the identification of secondary building blocks (see example of distinct L-T-P and L-T-U blocks detected on a set of PacBio reads).

**S1 Table.**
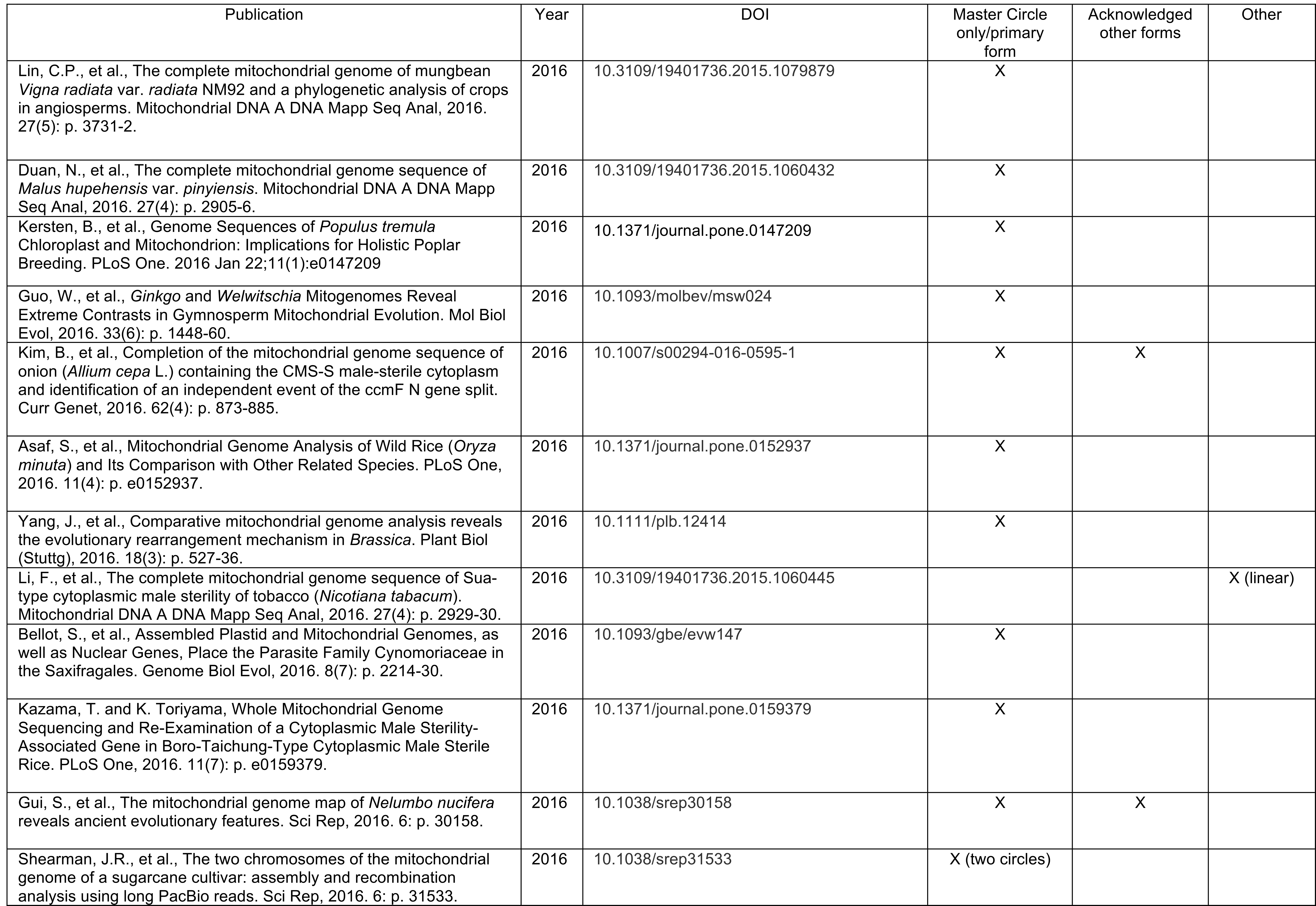


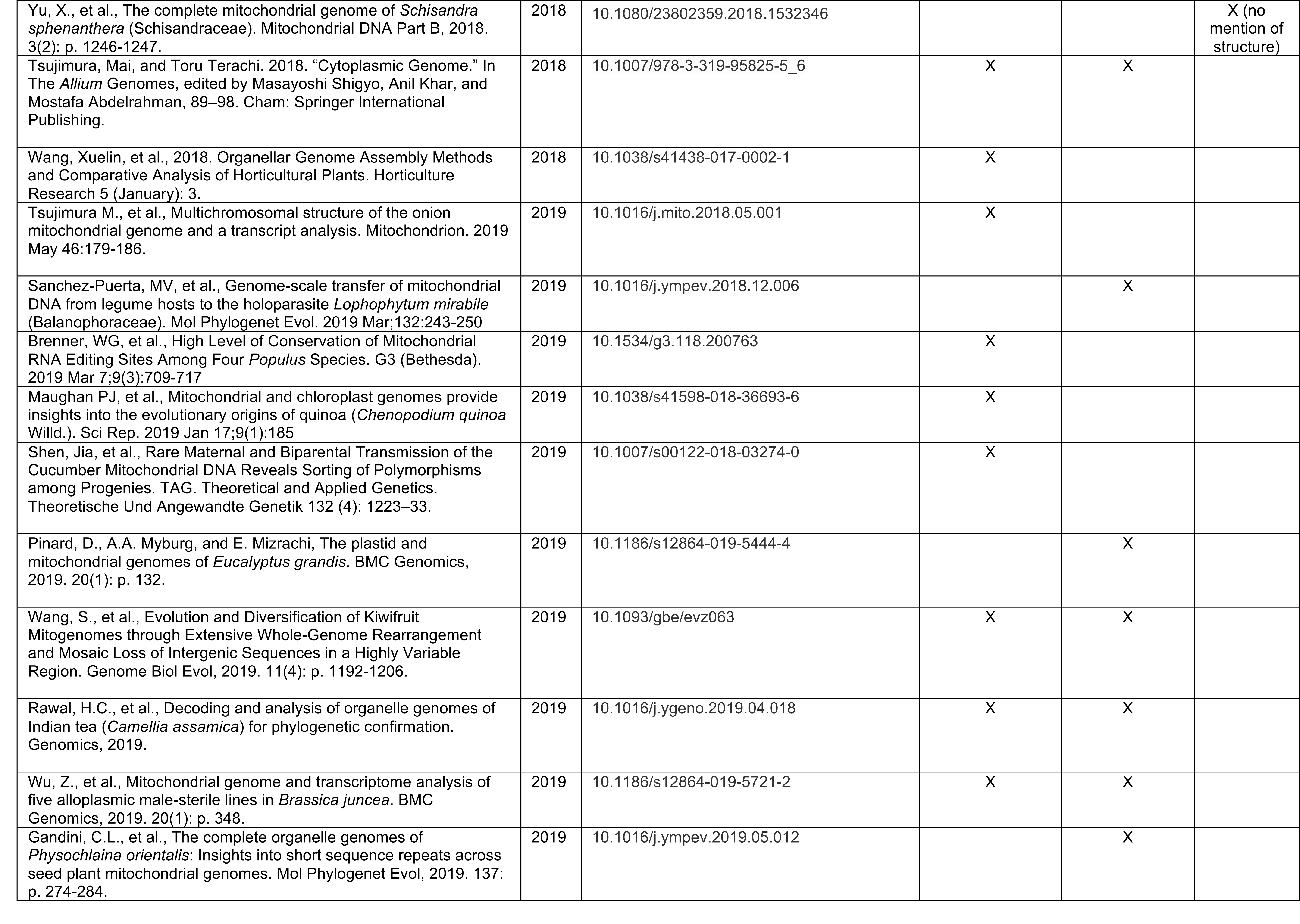
Plant mitochondrial genome publications from 2016–2019. A list of contemporary publications, many of which present a “master circle” model of the plant mitochondrial genome. The classifications reflect our best efforts to understand the data representation.

**S2 Table.** Structural mitochondrial genome composition of representative isoforms submitted to NCBI GenBank. Mitochondrial genome coordinates of primary structural units and recombinogenic medium size repeats on representative isoforms of corresponding GenBank files MK642355 (S2A), MK820672 (S2B), and MK759657 (S2C).

Mitochondrial genome coordinates of primary structural units on representative GenBank files S2A Table
| Primary structural units | Recombinogenic medium size repeats | Length | Orientation | *L.sativa* MK642355 (isoform A) |  |
| --- | --- | --- | --- | --- | --- |
|  |  |  |  | Start | End |
| N (copy #1) |  | 4116 | R | 1 | 4116 |
|  | X-01b (copy #1) | 576 | R | 3541 | 4116 |
| M (copy #1) |  | 30580 | R | 4117 | 34696 |
| K |  | 78272 | R | 34697 | 112968 |
| N (copy #2) |  | 4116 | R | 112969 | 117084 |
|  | X-01b (copy #2) | 576 | R | 116509 | 117084 |
| M (copy #2) |  | 30580 | R | 117085 | 147664 |
| Q |  | 11283 | R | 147665 | 158947 |
| T (copy #1) |  | 3552 | R | 158948 | 162499 |
| P |  | 20468 | R | 162500 | 182967 |
| R (copy #1) |  | 10430 | R | 182968 | 193397 |
| L |  | 44054 | R | 193398 | 237451 |
| T (copy #2) |  | 3552 | R | 237452 | 241003 |
| U |  | 38613 | R | 241004 | 279616 |
| R (copy #2) |  | 10430 | R | 279617 | 290046 |
| Z |  | 19295 | R | 290047 | 309341 |
|  | X-01b (copy #3) | 576 | F | 309342 | 309917 |
| W |  | 53983 | R | 309342 | 363324 |

S2B Table
| Primary structural units | Recombinogenic medium size repeats | Length | Orientation | *L.serriola* MK820672 (isoform B) |  |
| --- | --- | --- | --- | --- | --- |
|  |  |  |  | Start | End |
| N (copy #1) |  | 4116 | R | 1 | 4116 |
|  | X-01b (copy #1) | 576 | R | 3541 | 4116 |
| M (copy #1) |  | 30580 | R | 4117 | 34696 |
| K |  | 78272 | R | 34697 | 112968 |
| N (copy #2) |  | 4116 | R | 112969 | 117084 |
|  | X-01b (copy #1) | 576 | R | 116509 | 117084 |
| M (copy #2) |  | 30580 | R | 117085 | 147664 |
| Q |  | 11283 | R | 147665 | 158947 |
| T (copy #1) |  | 3552 | R | 158948 | 162499 |
| U |  | 38617 | R | 162500 | 201116 |
| R (copy #1) |  | 10430 | R | 201117 | 211546 |
| L |  | 44054 | R | 211547 | 255600 |
| T (copy #2) |  | 3552 | R | 255601 | 259152 |
| P |  | 20468 | R | 259153 | 279620 |
| R (copy #2) |  | 10430 | R | 279621 | 290050 |
| Z |  | 19295 | R | 290051 | 309345 |
|  | X-01b (copy #1) | 576 | F | 309346 | 309921 |
| W |  | 53983 | R | 309346 | 363328 |

S2C Table
| Primary structural units | Recombinogenic medium size repeats | Length | Orientation | *L.saligna* MK759657 |  |
| --- | --- | --- | --- | --- | --- |
|  |  |  |  | Start | End |
| M (copy #1) |  | 30534 | R | 1 | 30534 |
| K |  | 79109 | R | 30535 | 109643 |
| T (copy #1) |  | 4048 | F | 109644 | 113691 |
| L |  | 43583 | F | 113692 | 157274 |
| R (copy #1) |  | 10433 | F | 157275 | 167707 |
| P |  | 20420 | F | 167708 | 188127 |
| T (copy #2) |  | 4048 | F | 188128 | 192175 |
| Q |  | 11083 | F | 192176 | 203258 |
| M (copy #2) |  | 30534 | F | 203259 | 233792 |
|  | X-01a (copy #1) | 877 | F | 233793 | 234669 |
| UV |  | 90023 | R | 233793 | 323815 |
| R (copy #2) |  | 10433 | R | 323816 | 334248 |
| Z |  | 19278 | R | 334249 | 353526 |
|  | X-01a (copy #2) | 877 | F | 353527 | 354403 |
|  | S-01-IR (copy #1) | 1218 | F | 353527 | 354744 |
| S |  | 14743 | R | 353527 | 368269 |
|  | S-01-IR (copy #2) | 1218 | R | 367052 | 368269 |
|  | X-01a (copy #3) | 877 | R | 367393 | 368269 |

## Notes

#### Summary of Updates

Some new data added on structures of mitochondrial genomes, and some rewriting to reflect current thought in the field.

